# Quantitative proteomics and phosphoproteomics of PPP2R5D variants reveal deregulation of RPS6 phosphorylation through converging signaling cascades

**DOI:** 10.1101/2023.03.27.534397

**Authors:** KA Smolen, CM Papke, MR Swingle, A Musiyenko, C Li, AD Camp, RE Honkanen, AN Kettenbach

**Affiliations:** Department of Biochemistry and Cell Biology, Geisel School of Medicine at Dartmouth, Hanover, NH 03755, USA; Department of Biochemistry and Molecular Biology, University of South Alabama, Mobile, AL 36688, USA; Norris Cotton Cancer Center, Geisel School of Medicine at Dartmouth, Lebanon, NH 03756, USA

**Author notes:** Corresponding authors (R.E.H); (A.N.K).

**Keywords:** PPP2R5D, PP2A, Phosphatase, PPP2R5D-related neurodevelopmental disorder, Proteomics, Phosphoproteomics, mTOR

## Abstract

Variants in the phosphoprotein phosphatase-2 regulatory protein-5D gene (*PPP2R5D*) cause the clinical phenotype of Jordan’s Syndrome (PPP2R5D-related disorder), which includes intellectual disability, hypotonia, seizures, macrocephaly, autism spectrum disorder and delayed motor skill development. The disorder originates from *de novo* single nucleotide mutations, generating missense variants that act in a dominant manner. Pathogenic mutations altering 13 different amino acids have been identified, with the E198K variant accounting for ∼40% of reported cases. Here, we use CRISPR-PRIME genomic editing to introduce a transition (c.592G>A) in the *PPP2R5D* allele in a heterozygous manner in HEK293 cells, generating E198K-heterozygous lines to complement existing E420K variant lines. We generate global protein and phosphorylation profiles of wild-type, E198K, and E420K cell lines and find unique and shared changes between variants and wild-type cells in kinase- and phosphatase-controlled signaling cascades. As shared signaling alterations, we observed ribosomal protein S6 (RPS6) hyperphosphorylation, indicative of increased ribosomal protein S6-kinase activity. Rapamycin treatment suppressed RPS6 phosphorylation in both, suggesting activation of mTORC1. Intriguingly, our data suggest AKT-dependent (E420K) and -independent (E198K) activation of mTORC1. Thus, although upstream activation of mTORC1 differs between PPP2R5D-related disorder genotypes, treatment with rapamycin or a p70S6K inhibitor warrants further investigation as potential therapeutic strategies for patients.

## 1. INTRODUCTION

PPP2R5D-related intellectual disability (ID) and developmental delay (DD) disorder (OMIM#616355) is a syndrome characterized by mild to severe neurodevelopmental delay. The disorder, also known as Jordan’s Syndrome, is characterized by intellectual disability, seizures, epilepsy, macrocephaly, autism-spectrum-disorder (ASD), hypotonia, and delayed motor skill development (1–4). The PPP2R5D-related neurodevelopmental disorder is autosomal dominant, arising from *de novo* germline missense mutations in the *PPP2R5D* gene (1, 3, 5).

*PPP2R5D* encodes a regulatory subunit of type 2A serine/threonine phosphoprotein phosphatase (PP2A). Most PP2A phosphatases function as heterotrimeric holoenzymes, which are ubiquitously expressed in most cell types. To generate each PP2A-holoenzyme, a regulatory-targeting (B) subunit is assembled with a core dimer consisting of a scaffolding (A) subunit and a catalytic (C) subunit (6, 7) (**Figure 1A**). The A-C core dimer is expressed in most, if not all, human tissues, and humans express two highly similar isoforms of both the C and A subunits (*PPP2CA/ PPP2CB* and *PPP2R1A/PPP2R1B*, respectively). There are four families of B-subunits (B, B’, B’’, and B’’’/Striatin), and each family has several members (**Figure 1A**). Some B-subunits are widely expressed, while the expression of others is restricted to a subset of cell types (8). *PPP2R5D* encodes the delta isoform of the B’-family subunit and is called B’δ B56δ, PR61, and PPP2R5D in the literature. PPP2R5D/B56δ is ubiquitously expressed, with slightly higher levels reported in the brain, breast, testis, and gastrointestinal tissue (9–11). In addition to each B-family having several isoforms, additional isoforms are generated by alternate splicing (8, 12). Therefore, although commonly referred to as PP2A in the literature, combinatorially, the PP2A family may include over a hundred unique holoenzymes (**Figure 1A**).

**Figure 1.**
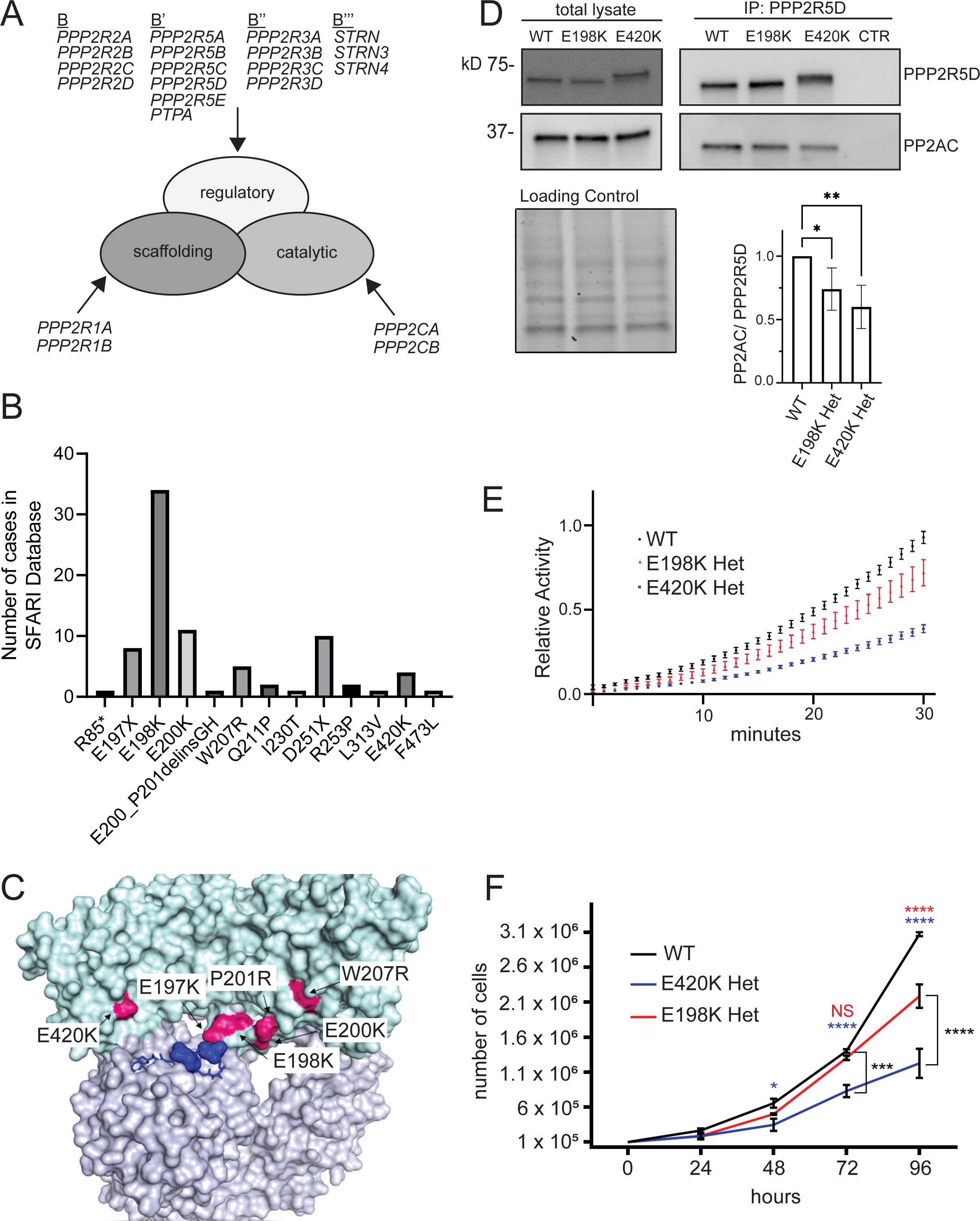
PP2A holoenzyme structure, PPP2R5D variants, and E198K vs E420K variant activity profiling. (A) Diagram of PP2A holoenzyme structure and components. (B) Number of cases corresponding to pathogenic PPP2R5D variants based on SFARI database. (C) PPP2R5D homology model showing the location of the common PPP2R5D variants seen in PPP2R5D-related neurodevelopmental disorder. The model is based on that of PPP2R5C (PDB ID: 2IAE), where PPP2R5D is colored in cyan and the scaffolding and catalytic subunits are gray. Metal ions are shown in dark blue. Pink corresponds to variant amino acids such as E198K and E420K. (D) Western analysis of total cell lysates (input) and endogenous PPP2R5D IPs probed for PPP2R5D or for the PP2A catalytic subunit (PP2AC). Relative amounts of PP2AC from WT, E198K-het, or E420K-het cells normalized to total PPP2R5D in IPs and represented as the mean percent of WT. (N=5 independent biological replicates, mean +/−SD, ANOVE Dunnett_’_s multiple comparison posthoc test *P<0.05, **<0.01) (E) Relative phosphatase activity of endogenous PPP2R5D IPs. Endogenous PPP2R5D was immunoprecipitated from WT, E198K-het, and E420K-het cells, and hydrolysis of DiFMUP was measured at Ex 360nm and at Em 460nm. Activity was normalized to total levels of PPP2R5D detected in IPs through immunoblotting shown in Figure 1D. WT activity (black) was normalized at 30 minutes (endpoint) to 1 and variant activity (E198K-het=red and E420K-het=blue) is shown relative to WT (n = 5 independent biological replicates of cells, each performed as n = 2 technical replicates). (F) Proliferation Assay of WT (black), E198K (red), and E420K (blue) variant cells. (N=3 independent biological replicates, each performed as n=2 technical replicates). Error bars represent the mean ± SD, two-way ANOVA Tukey’s, **** P<0.0001, *** P<0.001, ** P<0.01, * P<0.05, NS: non-significant).

To date, over 20 germline mutations in the *PPP2R5D* coding region have been reported, generating pathogenic missense mutations of 13 amino acids (13) (**Figure 1B, Supplemental Table 1**). Many pathogenic variants are charge reversal changes, in which a negatively charged acidic amino acid (e.g. glutamic acid, E) is mutated to a positively charged basic amino acid (e.g. lysine or arginine, K or R) due to a single base genomic alteration (e.g., glutamate (E), encoded by <u>G</u>AA or <u>G</u>AG, is converted to lysine (K), encoded by <u>A</u>AA or <u>A</u>AG). The most frequently observed pathogenic changes are p.Glu198Lys (E198K), p.Glu200Lys (E200K), p.Glu420Lys (E420K), and p.Asp251X (D251X, with X representing Ala, His, Val, or Tyr). Patients with E198K or E420K variants often display severe clinical symptoms (1, 14).

The E198 residue, its surrounding region, and much of the protein core are conserved between PPP2R5C/B56γ and PPP2R5D/B56δ (**Supplemental Figure 1**), and the crystal structure of a trimeric PP2A holoenzyme containing PPP2R5C (B56γ) has been solved (1, 15–17). In the PPP2R5C structure, the amino acids corresponding to E197, E198, and E200 in PPP2R5D reside in an acidic loop between α-helices 3 and 4. This loop is in close contact with the catalytic metals contained in the C-subunit of the PP2A-core dimer (1, 15–17), suggesting a role in the regulation of catalytic activity (**Figure 1C**). The prevalence of pathogenic variants in the corresponding loop in PPP2R5D suggests that the acidic loop is important for holoenzyme function (**Figures 1B**, **1C**). Homology models also suggest that other pathogenic variants (e.g. E197K, E200K, P201R, W207R, and E420K) are positioned on the surface, facing the active site of the catalytic subunit (**Figure 1C**) (1, 2, 14). The greater severity of symptoms observed in patients with the E198K variant compared to patients with mutations of E197 and E200 may also suggest that E198 has a more significant role in PP2A function (1, 2, 14). Previous biochemical studies have reported that when expressed *in vitro* as a tagged fusion protein in cells, the PPP2R5D E198K variant has a longer half-life than the wild-type (WT) (1). In addition, overexpression of PPP2R5D E198K in HEK293T cells increases the phosphorylation of PP2A-B56δ substrates, such as serine 9 of GSK-3β (1). These observations are consistent with dominant-negative suppression of PPP2R5D-dependent PP2A activities (1).

To decipher the molecular events associated with clinically relevant *PPP2R5D* variants, we adapted genomic editing methods to establish human cell lines recapitulating known pathogenic variants in PPP2R5D, allowing the study of how the expression and actions of endogenous variant proteins alter normal biological processes. Employing a fourth-generation single-base editing system (BE4-Gam), we previously generated heterozygous E420K variant cell lines (18). We reported that the endogenous E420K variant in HEK293 is associated with increased phosphorylation levels of several PPP2R5D substrates (18). However, the most common pathogenic variant (E198K) could not be generated using BE4-Gam without also causing non-synonymous bystander mutations, likely because BE4-Gam has a multi-base editing window and the desired c.592G>A mutation is located in a region surrounded by multiple cytosines encoding E197, D199, and E200. To overcome this technical challenge, we adapted a CRISPR-PRIME system, which utilizes a catalytically impaired Cas9 nickase (H840A mutant) fused to reverse transcriptase. CRISPR-PRIME is programmed with a pegRNA (<u>p</u>rime <u>e</u>diting <u>g</u>uide RNA) that specifies the target site and encodes the desired edit (**Supplemental Figures 2 and 3**). The methods described herein proved useful for introducing the desired nucleotide change (c.592G>A: p.E198K) in the genome of HEK293 cells and can likely be adapted to generate cell lines with almost any single base change.

**Figure 2.**
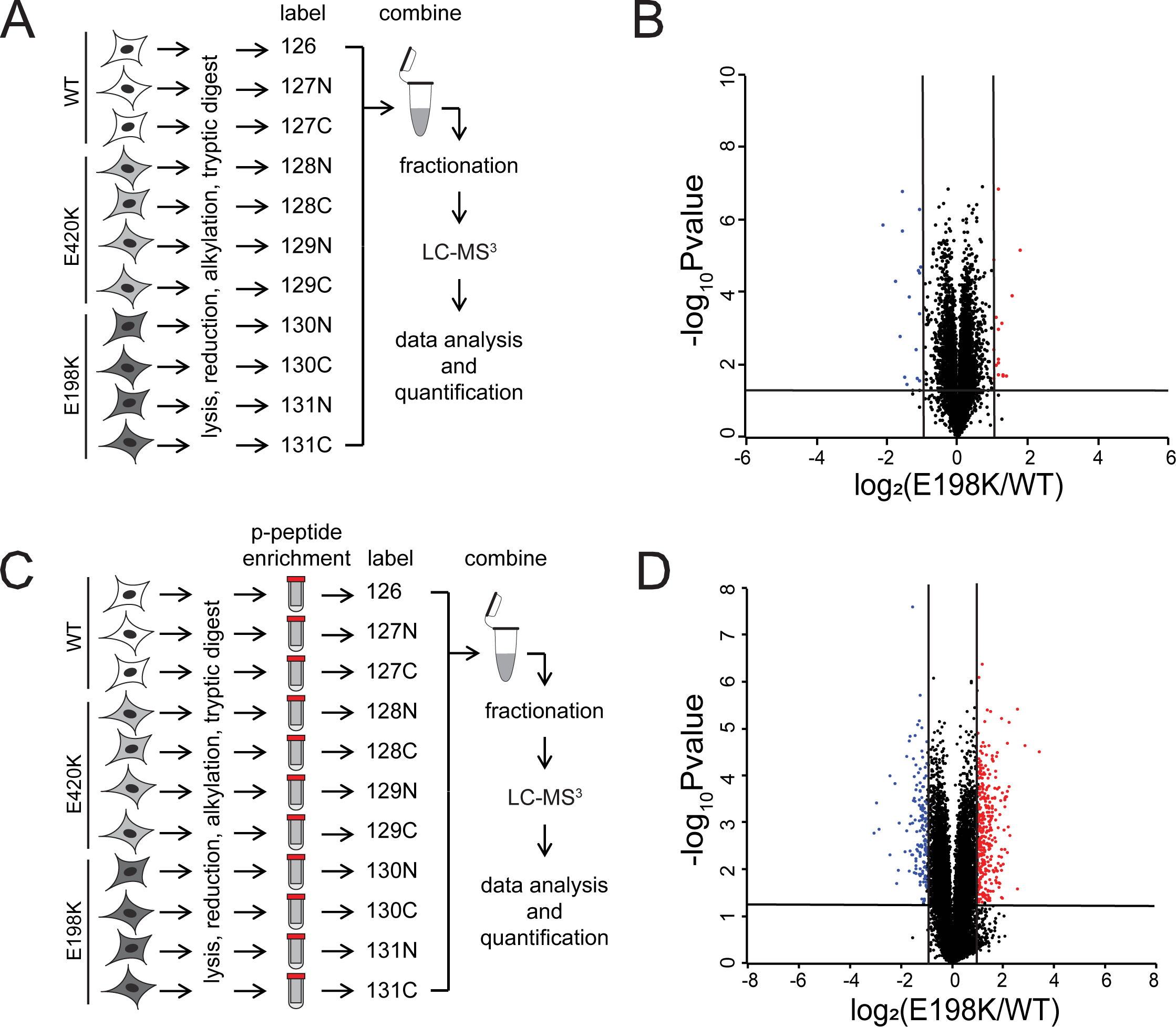
Quantitative changes in the proteome and phosphoproteome in PPP2R5D E198K variant cells. Workflow for the quantitative proteomics (A) and phosphoproteomics (C) protocol used. Volcano plots for proteomic (B) and protein corrected phosphoproteomic changes (D) in E198K variant cells (N=3 and N=4 independent biological replicates for WT and E198K-het, respectively.) Volcano plots show log_2_ fold change versus the negative log_10_ of the *p-*value of the fold change. Statistical significance corresponding to a *p*-value of <0.05 is shown by peptides with a -log_10_ value of 1.3 or greater. Peptides shown in blue or red are two-fold or more decreased or increased in abundance, respectively.

**Figure 3.**
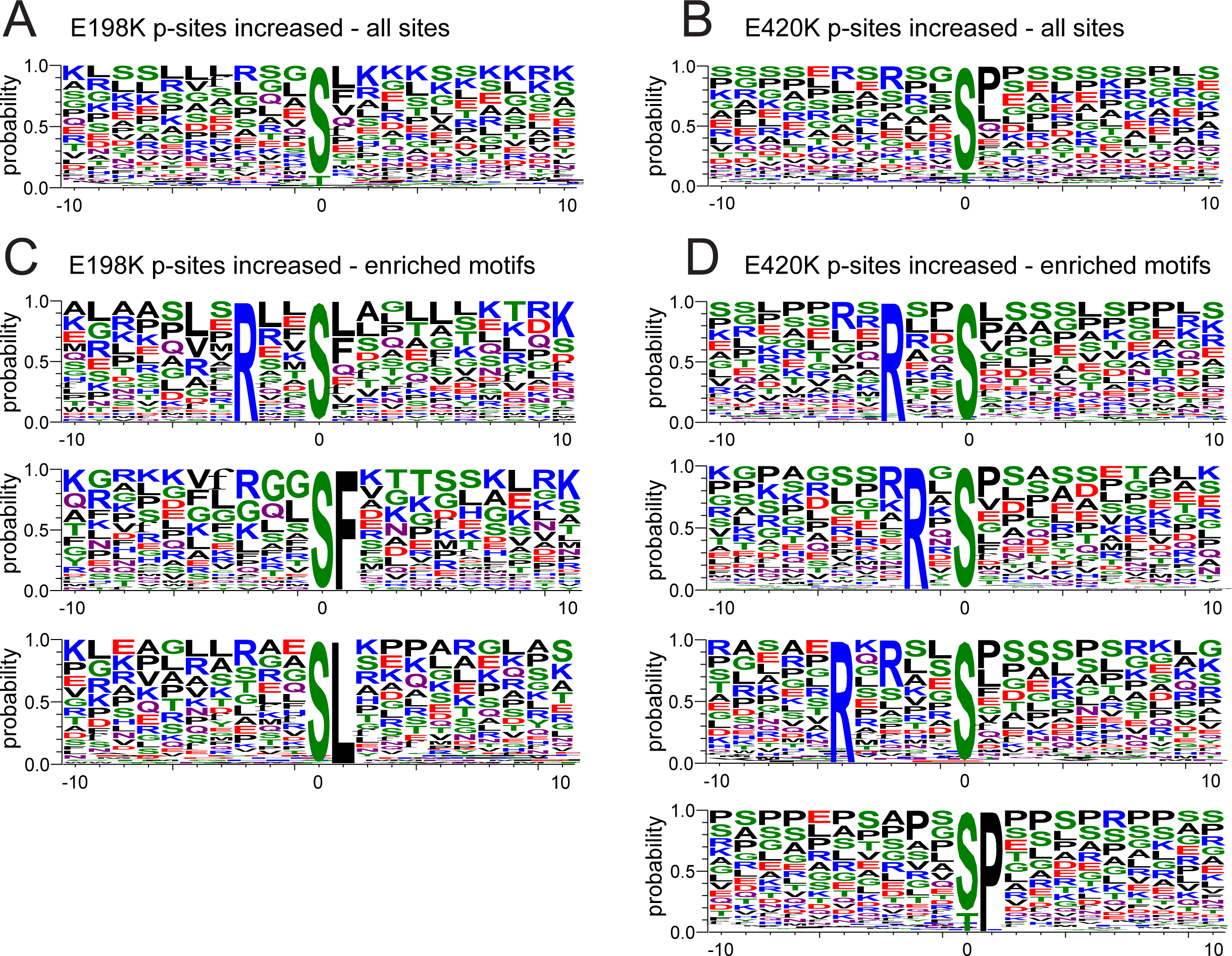
Phosphorylation site motif enrichment analysis of sites that were significantly increased in the E198K and E420K variant cell line. (A) Weblogos showing enrichment of amino acids surrounding the phosphorylated residue for sites significantly increased by at least two-fold in PPP2R5D E198K (A) and E420K (B) variant cells. (C) Enriched motifs among significantly increased phosphorylation sites in the E198K variant. (D) Enriched motifs among significantly increased phosphorylation sites in the E420K variant.

To investigate changes in PP2A-PPP2R5D function in E198K heterozygous cells, we performed quantitative proteomic and phosphoproteomic analyses to determine protein and phosphorylation abundance changes in WT, E198K, and E420K HEK293 cells. Our analyses reveal unique and common deregulated phosphorylation events in PPP2R5D E198K and E420K heterozygous cells. Intriguingly, we identify increased phosphorylation of an activating threonine (T389/T412) in p70 ribosomal protein S6 kinase (p70S6K/RPS6KB1/2), activating serines (S221 and S380) of p90 ribosomal protein S6 kinase 1 (RSK1/RPS6KA1), and (S232 and S389) of p90 ribosomal protein S6 kinase 6 (RSK4/RPS6KA6) with a corresponding increase in the phosphorylation of a common substrate (RPS6) as shared in both variant cell lines. Although the deregulation of the upstream regulatory mechanisms differs between the E198K and E420K variant cell lines, inhibition of mTORC1 with rapamycin or of p70S6K with LY2584702 reduces RPS6 phosphorylation in both variant cell lines. These results suggest inappropriate mTORC1 activation may be a common deregulation in PPP2R5D-related neurodevelopmental disorders. Thus, further investigation with rapamycin analogs, such as Everolimus, which has proven safe and effective in treating neuropsychiatric disorders (e.g. tuberous sclerosis complex syndromes (TSC), as well as LY2584702 treatment, which has been shown helpful to patients with ASD and fragile X, may be warranted to evaluate their potential as therapeutic options for patients with the most common PPP2R5D variants (19–21).

## 2. RESULTS

### 2.1. Generation of *PPP2R5D* E198K variant cell lines

To more broadly investigate signaling changes in the most common pathogenic PPP2R5D variant, we generated E198K variant cell lines. Our previous studies revealed alterations in phosphorylation-regulated signaling cascades in the PPP2R5D E420K variant. However, it is unclear if the different variants alter similar or distinct signaling pathways (18). To address this key question, we employed a PE3b, a third-generation prime editor (Anzalone et al., 2019) (**Supplemental Figures 2 and 3**) to introduce the pathogenic mutation (c.592G>A) into the genome of HEK293 cells, generating lines with the most common pathogenic variant (E198K). The PE3b system is a nicking Cas9 (nCas9) fused to a reverse transcriptase that is programmed by a synthetic prime editing guide RNA (pegRNA) that targets the Cas9 protein to the genomic locus and encodes the desired edit (**Supplemental Figure 2**). We designed a pegRNA that positioned the nCas9 to nick the to-be-edited strand upstream of the target site (pegRNA nick in **Supplemental Figure 2**). After electroporation with the PE3b-PRIME system, the cells were grown for 48 hours to ensure cell replication had occurred, which made the mutation permanent in the genome. The cells were then single-cell sorted, and the emerging colonies were screened for the desired c.592G>A base change by Sanger sequencing (**Supplemental Figure 3**). To ensure the clonal cell lines with the desired mutation were homogenous, each confirmed E198K line was single-cell sorted two additional times before further analysis. Complete genomic exon sequencing of parental E198K-het and E420K-het variant lines was performed to detect potential off-target editing and spontaneous somatic mutations that could have occurred during repeated single-cell sorting, as previously described (18). Cell lines with off-target or enriched somatic mutations in protein-coding regions were discarded.

### 2.2. Expression, holoenzyme assembly, activity, and proliferation of the PPP2R5D E198K variant

Western blot analysis of total lysates of the parental WT, the E198K clone, and a previously characterized heterozygous E420K clone (18) showed no difference in the level of endogenous PPP2R5D or PP2AC (catalytic subunit) expression (**Figure 1D**). To determine how the expression of PPP2R5D variants affects PP2A holoenzyme assembly and stability, we immunoprecipitated endogenous PPP2R5D and probed the precipitates for both PPP2R5D and PP2AC (**Figure 1D**). We found that in cells expressing PPP2R5D E198K, the amount of catalytic subunit that co-IPs with PPP2R5D was reduced by 25.97±7.5% (**Figure 1D**). For E420K, co-IP of the catalytic subunit was reduced by 56.1±1.1 %.

Next, we compared the phosphatase activity of PP2A holoenzymes containing WT or variant PPP2R5D (**Figure 1E**) using an established fluorescent assay that employs DiFMUP as a substrate (22–25). We immunoprecipitated endogenous PPP2R5D using a surplus of a previously characterized PPP2R5D-specific antibody that recognizes a near C-terminal epitope unique to PPP2R5D (18), and observed dephosphorylation of DiFMUP in all samples. When normalized to the amount of PPP2R5D immunoprecipitated, less activity was observed for both PPP2R5D variants. These results are consistent with the reduced levels of catalytic subunit in PPP2R5D IPs generated from the variant cell lines. When normalized to the amount of PP2A catalytic subunit detected in the IPs, the phosphatase activity was similar between WT, E198K, and E420K samples. This data is consistent with the reduction in phosphatase activity observed in the variants arising from reduced retention or reduced incorporation of PP2AC into PPP2R5D variant holoenzymes.

We also determined that PPP2R5D variants alter cell growth using a cell proliferation assay in which cells were plated at equal density, and growth curves were documented over four days. When compared to WT HEK293 cells, we observed slower rates of growth in cells heterozygous for PPP2R5D E198K or E420K, with a significant reduction in growth in E420K and E198K variant cells observed by 48 hours and 96 hours, respectively (**Figure 1F**).

### 2.3. Quantitative changes in the proteome and phosphoproteome of PPP2R5D E198K variant cells

To obtain an unbiased assessment of how the E198K and E420K PPP2R5D variants affect cellular signaling, we employed quantitative proteomics and phosphoproteomics using the Tandem-Mass-Tag (TMT)-MS^3^ approach as reported previously (18). HEK293 WT, E198K, and E420K variant cells were grown as asynchronous populations, collected, lysed, and digested into peptides with trypsin to analyze protein abundance (**Figure 2A**). To distinguish peptides from different cell lines and replicates, peptides were bar-coded by labeling with TMT reagent. After labeling, peptides were combined, fractionated, and analyzed by LC-MS^3^ (**Figure 2A**). In this analysis, we identified and quantified 8830 proteins, of which only 34 proteins (0.4%) were significantly increased or decreased at least two-fold in E198K variant cells compared to WT (**Figure 2B** and **Supplemental Table 2**). Consistent with our previous observations, only 312 (3.5%) proteins were significantly increased or decreased at least two-fold in E420K variant cells compared to WT (**Supplemental Figures 4A and 4B**). Importantly, we did not observe significant changes in the abundance of PP2A catalytic, scaffolding, and regulatory subunits in either PPP2R5D variant cell line (**Supplemental Table 3**).

**Figure 4.**
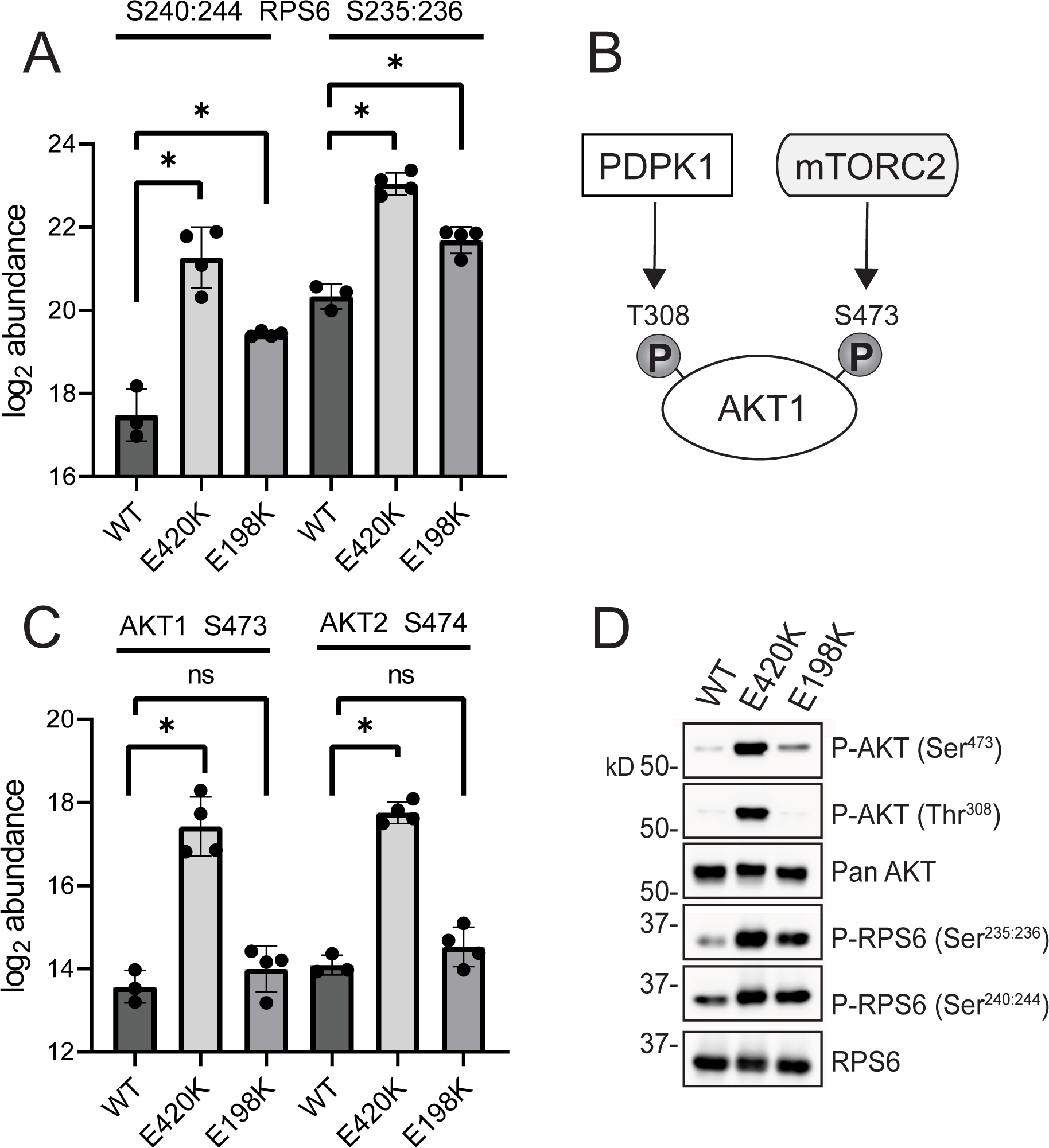
PPP2R5D variants display aberrant ribosomal protein S6 (RPS6) phosphorylation. (A) LC-MS^3^ data for sites S240:244 and S235:236 of RPS6 in WT, PPP2R5D E420K-het, and PPP2R5D E198K-het HEK293 cell lines. Value is the log_2_ of the TMT intensities detected via LC-MS^3^. (B) Activating phosphorylations of AKT serine-threonine protein Kinase 1 (AKT1) by 3-phosphoinsotide dependent protein kinase 1 (PDPK1) and mechanistic target of rapamycin kinase complex 2 (mTORC2). (C) Log_2_ LC-MS^3^ TMT intensities of S473 of AKT1 and S474 of AKT2 in WT, PPP2R5D E420K-het, and PPP2R5D E198K-het HEK293 cell lines. (D) Representative immunoblots showing the levels of P-AKT1/2 at S473/4 and T308/9, of P-RPS6 at S235:236 and S240:244, total AKT protein (Pan AKT) and total RPS6 protein in wild type (WT), PPP2R5D E420K-het, and PPP2R5D E198K-het HEK293 cell lysates.

To identify changes in phosphorylation abundances, we performed phosphopeptide enrichment, TMT labeled the phosphopeptides, identified the proteins from which the peptides were derived, and quantified the phosphopeptide intensities (**Figure 2C**). In this analysis, we identified and quantified 31,714 phosphorylation sites (**Supplemental Table 2**). Although we did not observe large changes in protein abundance, we corrected phosphopeptides for changes in the abundance of the respective protein. We identified and quantified the corresponding protein for 28,892 of the 31,714 phosphorylation sites, allowing us to perform protein correction on most phosphopeptides (**Supplemental Table 2**). After protein correction, we observed 612 (2.1%) phosphorylation sites that were significantly increased or decreased by at least a two-fold log_2_ change for the E198K variant (**Figure 2D**, **Supplemental Table 2**). The majority of these sites, 70%, were increased, and 30% were decreased. For the E420K variant, we identified 1751 (6.1%) phosphorylation sites to be significantly different, of which 78% were increased in phosphorylation (**Supplemental Figure 4D, Supplemental Table 2**). The higher number of phosphorylation sites with an increase in occupancy in E420K versus E198K variant cells is consistent with the greater decrease in catalytic subunit association and enzymatic activity that was observed for the E420K variant PP2A holoenzyme, measured using immunoprecipitation and *in vitro* phosphatase activity analysis, respectively (**Figure 1E**).

### 2.4. Alterations in phosphorylation signaling in PPP2R5D variant cells

To determine which signaling pathways are altered in the variant cells, we investigated the nature of fully-localized, singly-phosphorylated sites. We predicted the kinases that might be responsible for their phosphorylation. First, we performed phosphorylation site motif enrichment analysis of sites that were significantly increased (**Figures 3A and B**) or decreased (**Supplemental Figure 5A and 5B**) by two-fold or more in each variant. For sites that increased in the E198K variant cells, we observed an enrichment of arginine in the -3 position and phenylalanine or leucine in the +1 (**Figure 3C**). We found the same arginine enrichment in the -3 position for E420K variant cells and a preference for arginine in the -5 or proline in the +1 position (**Figure 3D**). Phosphorylation sites that significantly decreased were characterized by acidic amino acids downstream of the phosphorylation site in both variants (**Supplemental Figure 5A and 5B**).

**Figure 5.**
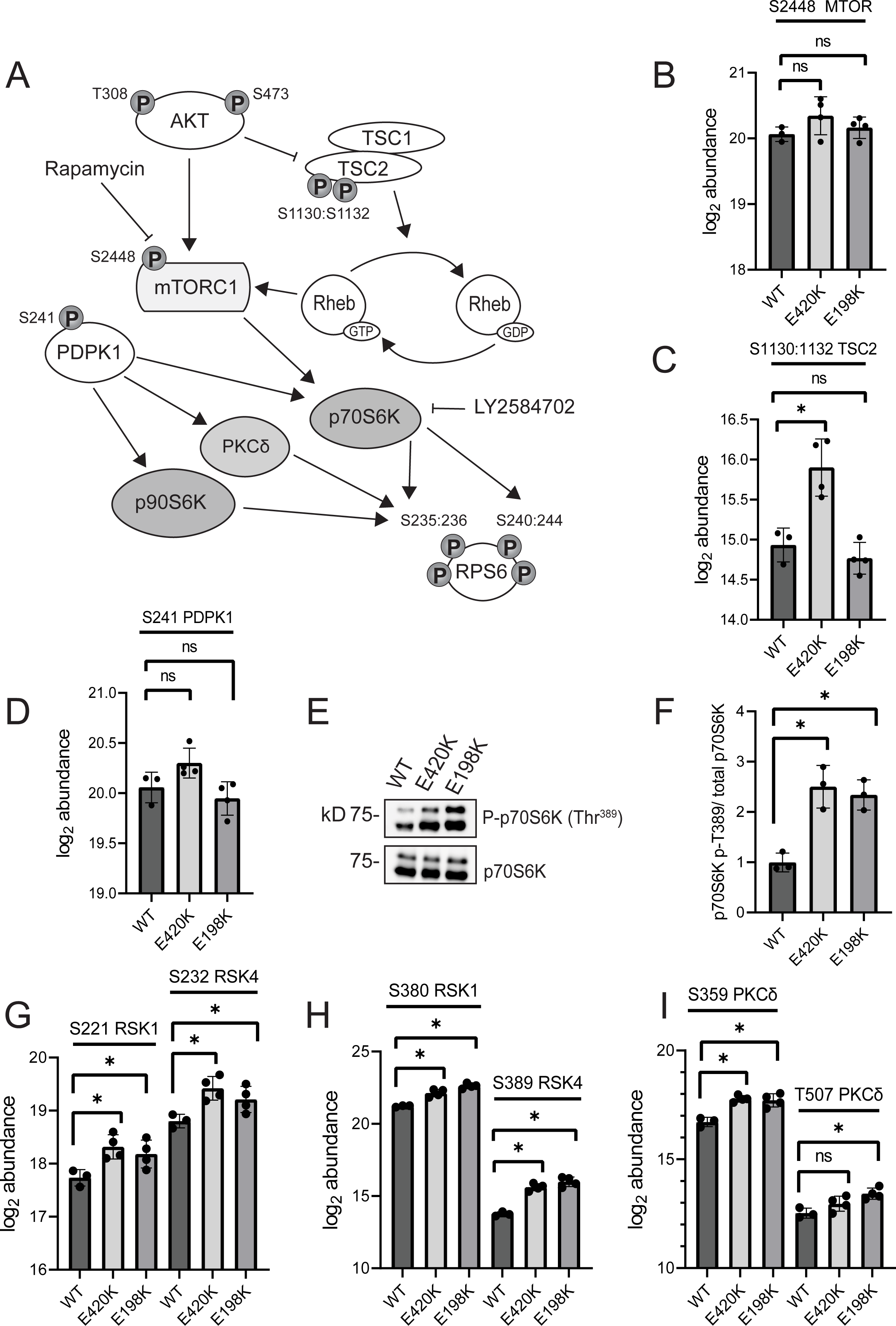
Phosphorylation changes in AKT-mTORC-RPS6 signaling cascades. (A) Diagram of the AKT-mTOR-RPS6 pathway. LC-MS^3^ expression data in WT, PPP2R5D E420K-het, and PPP2R5D E198K-het HEK293 cell lines is shown for phosphorylation of S2448 of mTOR (B), S1130:1132 of TSC2 (C), and S241 of PDPK1 (D). Values are log_2_ TMT intensities detected via LC-MS^3^. (E) Representative immunoblot showing the level of P-p70S6K T389 in WT, PPP2R5D E420K-het, and PPP2R5D E198K-het HEK293 cell lines. (F) Associated graph of P-T389 of p70S6K normalized to total p70S6K. LC-MS^3^ TMT intensities in WT, PPP2R5D E420K-het, and PPP2R5D E198K-het HEK293 cell lines is shown for phosphorylation of S221 of RSK1 and S232 of RSK4 (G), S380 of RSK1 and S389 of RSK4 (H), and S359 and T507 of PKCδ. Values are log_2_ TMT intensities detected via LC-MS^3^.

To learn more about the kinases that phosphorylate these sites, we used KinomeXplorer to predict upstream kinases of the regulated phosphorylation sites (26). Analysis of phosphorylation sites that increased in variant cells revealed a prevalence of Protein Kinase C (PKC)-dependent phosphorylation sites in both the E198K and E420K variant cells (**Supplemental Table 4**), which is consistent with an enrichment of arginines in the -3 and larger hydrophobics (leucine or phenylalanine) in the +1 position (**Figure 3C** and **3D**). In E420K variant cells, we also observed protein kinase B-(AKT-) and extracellular signal-regulated kinase (ERK) 1/2-dependent motifs and enrichment of consensus sequences of proline-directed kinases, such as cyclin-dependent kinases (CDKs), glycogen synthase kinase-3 (GSK3), and mitogen-activated protein kinases (MAPKs) (**Figure 3D**) (27). Of note, the enrichment of phosphorylation sites with arginine in the -3 and -5 positions (RxRxxS/T) is consistent with the phosphorylation site preference of AKT. This enrichment was previously observed in our studies of E420K variant HEK293 cells (18) but was not apparent in the E198K data. For phosphorylation sites that decreased, we detected a preference for serine-directed sites with proline in the +1 position and the enrichment of acidic amino acids (e.g. E, D) downstream of the phosphorylation site in both datasets. This may indicate deregulation of casein kinase 2 (CK2) in the variants (**Supplemental Table 4**) (27).

### 2.5. PPP2R5D variants display aberrant ribosomal protein S6 (RPS6) phosphorylation

Studies using inhibitors (e.g. okadaic acid) of the common catalytic subunit (C subunit) or siRNA targeting the common core scaffold (A subunit) have revealed that, as a family, PP2A-holoenzymes influence nearly all signaling cascades controlled by reversible phosphorylation (8). Previously, we found that E420K variant cells are characterized by altered insulin-, mTOR-, AMP-activated protein kinase (AMPK)-EGF receptor family (ErbB)-signaling, and additional downstream effectors. Notably, in the E420K variant, we observed constitutively active AKT-signaling associated with hyperphosphorylation of the downstream protein RPS6 (18). Our phosphoproteomic data confirmed hyperphosphorylation of RPS6 on S235:236 and S240:244 in E420K and revealed a similar increase in E198K variant cells (**Figure 4A**). However, although we found an enrichment of an AKT consensus motif (RxRxxS/T) in phosphorylation sites with significantly increased abundance in E420K cells, we did not observe this motif in phosphorylation sites increased in the E198K variant (**Figure 3C**). This prompted us to interrogate the AKT signaling pathway further. Human cells express three isoforms of AKT (AKT1/2/3), and their kinase activity is activated by phosphorylation at T308/9/5 and S473/4/2 by 3-phosphoinositide dependent protein kinase 1 (PDPK1) and mTORC2, respectively (**Figure 4B**) (28, 29). In our phosphoproteomic data, the amino acid sequence of phosphopeptides allows us to distinguish the AKT isoforms. Compared to WT, we observed an increase in AKT1/2 phosphorylation at S473/4 in E420K but not E198K variant cells, which is consistent with the results of the motif analysis (**Figure 4C**). For orthogonal validation of this observation, we performed western blot analysis of WT, E420K, and E198K variant cells using site-specific phospho-antibodies (**Figure 4D**) and observed an increase in AKT1 S473 phosphorylation in E420K but not in E198K variant cells compared to WT (**Figure 4D**). Western analysis also revealed increased phosphorylation of the activating T308 site in E420K variant cells (**Figure 4D**). Both AKT1 T308 and S473 phosphorylation was increased by greater than 20-fold in the E420K variant cells as compared to WT or E198K. These data suggest that the increase in RSP6 phosphorylation occurs in an AKT-dependent manner in the E420K variant and is independent of AKT in the E198K variant.

### 2.6. Activation status of kinases downstream of AKT

To identify the pathway resulting in increased RPS6 phosphorylation in E198K variant cells, we further explored the AKT-mTORC1 pathway by analyzing activating phosphorylations of downstream AKT targets (**Figure 5A**). Upon activation, AKT phosphorylates and activates mTOR on S2448 when mTOR is part of the mTORC1 complex (30–32). However, the phosphoproteomic analysis did not reveal a significant increase in this phosphorylation site in either variant (**Figure 5B**). mTORC1 activity is also regulated by the small GTPase Rheb, and the tuberous sclerosis complex (TSC) subunits 1 and 2 act as a dimer to negatively regulate Rheb-mediated mTORC1 activation (33, 34). AKT phosphorylation of TSC2 at S1130 and S1132 inactivates the TSC1/TSC2 complex (33, 35). We found that TSC2 phosphorylation at S1130 and S1132 is increased in E420K but not E198K variant cells (**Figure 5C**). These data are consistent with the activation of mTORC1 in E420K variant cells via an indirect mechanism in which altered AKT-dependent phosphorylation of TSC2 leads to the inactivation of the inhibitory TSC1/2 complex.

RPS6 is phosphorylated by a family of ribosomal protein S6 kinases (S6Ks) and protein kinase C delta (PKCδ) (36, 37). S6K-family members are activated by “priming” phosphorylation(s) in concert with activating T-loop phosphorylation(s) (38, 39) to release autoinhibition and realign active-site residues to capture the substrate and catalyze phosphorylation. There are two subfamilies of ribosomal protein S6 kinases: p70 ribosomal protein S6 kinases (p70S6Ks: p70S6K-alpha= RPS6KB1 and p70S6K-beta= RPS6KB2) and the p90 ribosomal protein S6 kinases (p90S6Ks: RPS6KA1=RSK1, RPS6KA2=RSK3, RPS6KA3=RSK2 and RPS6KA6 *=* RSK4). p70S6Ks are primarily activated via the AKT-mTORC1 pathway (40, 41). Interestingly, we found a significant increase in activating p70S6K phosphorylation (T389) in both the E420K and E198K variants, a site known to be phosphorylated by mTORC1 (40, 41) (**Figure 5E** and **5F**). The p90S6Ks, also referred to as ribosomal S6 kinases (RSKs), are activated via the mitogen-activated protein kinase (MAPK) pathway and 3-phosphoinositide-dependent protein kinase 1 (PDPK1) (39, 42). We did not observe increased phosphorylation of S241 on PDPK1, which is believed necessary for activation (43) (**Figure 5D**). However, both activating phosphorylations on RSK1 (S221 T-loop, S380 linker) and RSK4 (S232 T-loop, S389 linker) are increased in both the E198K and E420K variants (**Figure 5G** and **5H**). We also observed increased phosphorylation of two known activating sites on PKCδ: S359 and T507 (**Figure 5I**). S359 of PKCδ is an activating phosphorylation site occurring in response to oxidative stress; this site was significantly increased in both the E420K and E198K variants (44). T507 is phosphorylated by PDPK1 (45) and was only significantly increased in the E198K variant. Taken together, although we only observed activating phosphorylation of AKT in E420K variant cells, downstream kinases responsible for RPS6 phosphorylation, including p70S6K, p90S6Ks/RSKs, and PKCδ, are all phosphorylated on sites known to activate kinase activity.

### 2.7. Inhibition of RPS6 phosphorylation in variant cells

In both variant cell lines, we observed increased p70S6K T389 phosphorylation, a direct substrate of mTORC1, and its downstream target RPS6 (S240:244). However, only E420K variant cells displayed hyperactivation of AKT, which is known as a principle mTORC1 activating kinase. Thus, to determine if RPS6 phosphorylation would respond to upstream kinase inhibition in both variant cell lines, we treated WT and variant cells with rapamycin (10 nM for 1 hour), an mTORC1 inhibitor, or LY2584702 (20 nM for 3 hours), a p70S6K inhibitor. We observed that the hyperphosphorylation of p70S6K T389 and RPS6 S240:244 was reduced in WT as well as the E420K and E198K variants by rapamycin treatment (**Figure 6A-C** and **Supplemental Figure 6**). Furthermore, inhibition of p70S6K with LY2584702 reduced RPS6 phosphorylation at S235:236 and S240:244 in the E420K and E198K variant cell lines (**Figure 6D-F**). Of note, after treatment with LY2584702, we observed an increase in p70S6K T389 phosphorylation in WT and variant cell lines and in AKT S473 phosphorylation in WT and E198K variant lines (**Supplemental Figure 7**). Previous reports also observed increased phosphorylation of p70S6K T389 upon LY2584702 treatment, which was explained by an increased association of mTOR with p70S6K (46). Thus, our studies indicate that, although AKT is only hyperphosphorylated at activating residues in E420K PPP2R5D variant cells, activation of mTORC1 and p70S6K signaling occurs in both E198K and E420K variant cells. Furthermore, the downstream aberrant RSP6 hyperphosphorylation is suppressed by either rapamycin or LY2584702 treatment.

**Figure 6.**
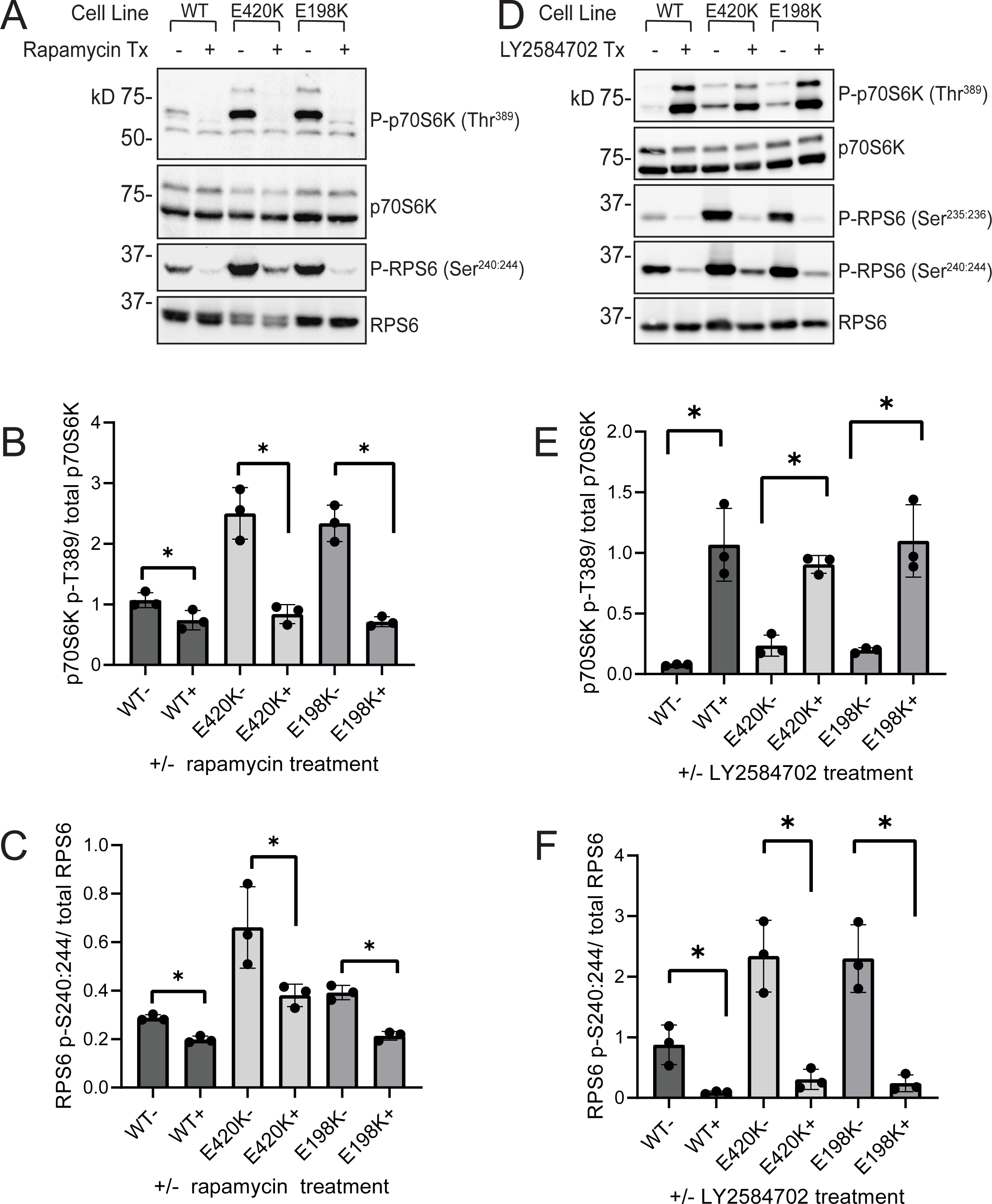
Reversal of aberrant mTOR signaling by rapamycin and p70S6K inhibitor (LY2584702) in E198K and E420K variant cells. (A and D) Representative immunoblot showing levels of P-p70S6K T389 and P-RPS6 in WT, PPP2R5D E420K-het, and PPP2R5D E198K-het HEK293 cell lines treated with rapamycin (10 nm for 1 hour) or LY2584702 (20 nM for 3 hours), a p70S6K inhibitor. Quantification of P-T389 of p70S6K normalized to total p70S6K for rapamycin treatment (B) and LY2584702 treatment (E). Quantification of P-S240:244 of RPS6 normalized to total RPS6 for rapamycin (C) and LY2584702 treatment (F). All treatments were performed as n=3 independent cell experiments, mean ± SD, unpaired *t*-test * *p* < 0.05.

## 3. DISCUSSION

Advances in DNA-sequencing technologies have enabled the identification of a plethora of novel pathogenic genomic variants, including over 20 spontaneous germline mutations in *PPP2R5D* recently linked to Jordan’s Syndrome (PPP2R5D-related neurodevelopmental disorder) (14). Additionally, pathogenic variants in PP2A-scaffolding, PP2A-catalytic, and other regulatory subunits of PP2A-holoenzymes have also been reported to cause neurodevelopmental disorders (47). Now the challenge is to determine the etiology associated with these genomic mutations and to develop interventions for the medical management of the associated disorders.

The phosphoprotein phosphatase 2A (PP2A) family is responsible for most phosphoserine and phosphothreonine dephosphorylation within a cell (8, 48, 49). Nonetheless, the roles of individual holoenzymes are poorly characterized. To determine the etiology of *PPP2R5D* variant disorders, we utilized a CRISPR-genomic single-base editing method that allowed the generation of HEK293 cell lines containing a heterozygous transition (c.1258G>A) in *PPP2R5D* on chromosome 6, precisely recapitulating a clinically relevant E420K variant. Characterization of the HEK-E420K cell lines revealed that the E420K variant altered a critical AKT-mTOR mediated signaling pathway that controlled cell growth (18). Here, we expanded this analysis to the most common *PPP2R5D* variant, E198K, to determine if the different variants alter the same, similar, or different biological processes.

The development of the E420K variant cell line was aided by the ideal position of a protospacer adjacent motif (PAM) that precisely positioned the single base editor to the target sequence. However, for targeting c.592G>A, which was needed to generate the E198K variant, this strategy failed, likely because of the additional cytosines were deaminated as bystander edits of the several nucleotide active site window of the system in the proximal genomic environment. To overcome this, we used the Prime Editor 3b (PE3b) strategy (50). The PE3b system is based upon a nicking Cas9 (nCas9) fused to a reverse transcriptase that is programmed by a synthetic prime editing guide RNA (pegRNA), which both targets the specific genomic locus and encodes the desired edit. With PE3b, the edit occurs without the need for double-stranded breaks or a donor template DNA for homology-directed repair. To facilitate prime editing, we designed a pegRNA that positioned the nCas9 to nick the to-be-edited strand upstream of the target site (pegRNA nick in **Supplemental Figure 2A**) and were successful in introducing the desired nucleotide change in the genome of HEK293 cells. To ensure the clonal cell lines with the desired mutation were homogenous, each confirmed E198K line was single-cell sorted two additional times prior to further analysis. Complete genomic exon sequencing of parental E198K-het and E420K-het variant lines was performed to detect off-target editing and spontaneous somatic mutations (Papke et al., 2021). This method should be easily adaptable for the generation of cell lines containing almost any novel single base or small region (1-4 bases) of DNA identified as a pathogenic genomic mutation.

For the study of variant PP2A-holoenzyme complexes, an advantage of employing genomic editing versus exogenous expression is that with genomic editing the expression, translation, and biogenesis of the variant holoenzymes are controlled by the endogenous cellular machinery. For PP2A holoenzymes, this is particularly important because their biogenesis involves the coordinated actions of many proteins (e.g., alpha4/IGBP1, PPME1, PTPA/PPP2R4, TIPRL, and LCMT1) (8, 51). During biogenesis, the common catalytic subunit is initially inactive. In a poorly understood process, it then undergoes several conformational changes and modifications before it is incorporated into the common A/C core dimer (52). PP2A-holoenzymes become functional after a unique B-subunit is integrated with the A/C core dimer. The proteins involved in the biogenesis of PPP2R5D holoenzymes are common for all PP2A holoenzymes found in most cells (12, 53). Thus, in studies where B-regulatory subunits are expressed exogenously, the interpretation of the data could be biased by the possibility that the biogenesis of other PP2A holoenzymes may be altered (e.g. overexpression of a B subunit may overload the biogenesis complex or sequester a limited number of A/C-subunits from other PP2A-holoenzymes).

The studies presented here revealed that the PPP2R5D WT and both E198K and E420K variants are expressed at similar levels. However, when PPP2R5D is immunoprecipitated, the IPs of both E198K and E420K cell lysates contained less catalytic subunit (PP2AC). This might be due to reduced stability of the variant holoenzymes, altered efficiency of the biogenesis process, altered “recycling” of variant-containing holoenzymes, or a pool of unbound variant PPP2R5D in the cells. This aspect of PPP2R5D biology warrants further investigation in future studies of the PPP2R5D-related neurodevelopmental disorder.

As previously reported for the E420K variant, the global proteome of the PPP2R5D E198K variant cells was stable, with only 0.4% of protein being either increased or decreased compared to WT. On the phosphorylation site level, we confirmed our findings related to the E420K variant, showing ∼ 6.1% of phosphopeptides were significantly increased or decreased by two-fold or more in the E420K variant cell line compared to WT. Interestingly, while the E198K also increases phosphopeptide occupancy in a heterozygous dominant fashion, the E198K variant had notably fewer (only 2.1%) significantly increased or decreased phosphopeptides. In both cell lines, the abundance of the catalytic, scaffolding, and regulatory subunits of PP2A were unchanged.

To identify potential therapeutic strategies for the management of PPP2R5D-related disorders, we interrogated signaling cascades altered by PPP2R5D variants. Previously, we found that the E420K PPP2R5D variant was associated with increased AKT-mTORC1 axis signaling leading to RPS6 hyperphosphorylation at S235:236 and S240:244 due to constitutive activation of AKT (18). Our results in this study were consistent with our prior observations related to altered AKT-activation in E420K based on increased T308 and S473 phosphorylation (28). In addition, our dataset clarifies a pathway by which mTORC1 is activated in the E420K variant cells. Notably, our data is not consistent with direct AKT-dependent phosphorylation of mTOR. Instead, we found that TSC2 phosphorylation at S1130 and S1132 was increased. AKT is known to phosphorylate these sites, which are known regulatory sites controlling the ability of TSC1 and TSC2 to act as a dimer in the negative regulation of Rheb-mediated mTORC1 activation (33, 35). Thus, our observations are consistent with mTORC1 activation in the E420K variant occurring by an indirect mechanism in which aberrant AKT-activation activates mTORC1 via phosphorylation of TSC2, which leads to the inactivation of the inhibitory TSC1/2 complex.

A major observation in this study was that the activating phosphorylations on AKT observed in the E420K variants do not occur in the E198K variant cells. However, increased phosphorylation of RPS6 on both S235:236 and S240:244 was observed in both variant cells. RPS6 is known to be phosphorylated at these sites by mTORC1-activated p70S6Ks (36, 54). Indeed, we detected hyperphosphorylation of p70S6K on T389 in both variants by western blot. RPS6 S235:236 can also be phosphorylated in a mTORC1-independent manner by p90S6Ks/RSKs and PKCδ (36, 55). We observed increased phosphorylation in both variants on RSK1 (S221 T-loop, S380 linker) and RSK4 (S232-T-loop, S389 linker). RSK1 S221 and RSK4 S232 are known targets of PDPK1 and phosphorylation of these residues is required for kinase activation (55). However, we did not observe increased activating phosphorylation of PDPK1 or altered phosphorylation of known regulatory residues on other established substrates for PDPK1 (e.g. RPS6KA3/RSK2, PRKACA, PRKCZ, PAK1, PKN1, or PKN2). This suggests that the variant phosphatases are not acting directly on PDPK1. We also observed increased phosphorylation of two known activating sites on PKCδ: T507 (up in the E198K variant) and S359 (up in both variants). At this time, the precise roles of PDPK1 and RPS6Ks in aberrant E198K and E420K variant signaling are not clear. Nonetheless, the ability of both rapamycin and LY2584702 to suppress aberrant RPS6 hyperphosphorylation suggests that further studies should be conducted to determine if other PPP2R5D variants display similar sensitivities and if these agents will prove helpful in the medical management of Jordan’s Syndrome.

Our observations linking E198K and E420K variants to altered mTOR signaling suggest Jordan’s syndrome should be added to a growing list of disorders associated with the dysregulation of mTOR pathways. Notably, dysregulation of mTOR has been implicated in Autism Spectrum Disorder (ASD) and other neurological and neurodegenerative disorders (30, 56, 57). Genetic syndromes, such as fragile X syndrome, tuberous sclerosis complex (TSC), PTEN-mediated ASD, neurofibromatosis (NF1) and others (56) are often heterogeneous at a clinical and genetic level, yet numerous studies have shown convergence of key biochemical pathways involving PI3K, AKT, and mTOR (56, 57). The PI3K/AKT/mTORC1 pathway is also known to regulate neuronal processes, including cell growth and survival, protein synthesis, excitatory synapse formation, and dendritic outgrowth (56). Deregulation of the pathway also results in symptoms such as epilepsy and megalencephaly, which are seen in Jordan’s Syndrome and other ASD (14, 56).

The development of PI3K/mTORC1/p70S6K inhibitors raises the possibility that specific targeting of this pathway may lead to therapeutic advances and the use of rapamycin and rapalogs has already been studied in the context of various pathologies. Rapamycin has been used for the management of multiple types of cancer, neurodegenerative diseases, including Alzheimer’s and Parkinson’s, and allograft rejection (19, 58, 59). In a mouse model of TSC, rapamycin reversed behavioral abnormalities and reduced epileptic activity (56, 57). However, the use of more specific drugs, such as those that target p70S6K or RSKs may be better treatment options due to the broad side-effects of mTOR inhibition (60, 61). In fragile X mice, seizures that are characterized by elevated RSK signaling can be prevented with RSK inhibitors (61). Several p70S6K inhibitors are in development or clinical trials, and the use of rapamycin or p70S6K inhibitors during early development of PTEN-ASD in mice was able to prevent the emergence ASD-like behavior (62). Similarly, rapamycin and p70S6K inhibitors have been shown to normalize dendritic spine density and reverse neurocognitive deficits in a mouse model of Angelman Syndrome (63, 64). In summary, our studies revealed AKT-dependent (E420K) and -independent (E198K) activation of mTORC1 and p70S6K as key signaling alterations in PPP2R5D-variant cells. We also observed increased activating phosphorylations on p90S6Ks (RSK1 and RSK4) and PKCδ with no apparent link to mTORC1 activation in both E198K and E420K variant cells. Although the affected pathways are complex and the variant PPP2R5D actions are not yet clearly defined, the pathways identified can be targeted with known drugs that inhibit mTORC1, such as rapamycin or rapamycin analogs and drugs that inhibit p70S6K (e.g. LY2584702). Based on our observations, these agents should be further tested to determine if they can restore normal signaling in a manner that will be beneficial. Exploring mTOR activation in the context of other PPP2R5D variants will help us to gain insight into their complex signaling and determine if targeting this pathway would be of clinical benefit to patients with PPP2R5D-related disorders.

## 4. METHODS

### 4.1 Cell Culture

HEK293 (Clontech #C3003-1; lot #7030396) cell lines were cultured in Dulbecco’s Modified Eagle’s Medium (DMEM), containing 25 mM glucose, 4 mM L-glutamine, 1 mM sodium pyruvate, 10 mM MEM non-essential amino acids, 100 units/mL penicillin, 100 μg/mL streptomycin (Gibco, Life Technologies), and 10% heat-inactivated Fetal Bovine Serum (Atlanta Biologicals, Lot# K19151) at 37°C with 5% CO2 in a humidified incubator. Cells were routinely passed at 70-90% confluency.

pCMV-PE2 (Addgene plasmid #132775; https:www.addgene.org/132775/) was provided as a generous gift from Dr. David Liu. This plasmid was amplified in E.coli DH5α and isolated using a Qiagen plasmid DNA purification kit (12123) according to the manufacturer’s protocol and was sequence verified. A dual RNA expression cassette containing sequences for the pegRNA (driven by an hU6 promoter) and the secondary nicking sgRNA (driven by a 7SK promoter) was synthesized, subcloned into pUC57-kan, and sequence verified by GenScript (Piscataway, NJ), which also performed plasmid preparation services to provide transfection grade DNA. In addition to the common sgRNA scaffold sequence, the pegRNA incorporated a target (protospacer) sequence (AGGGGCTGAGTTTGACCCAG), a primer binding site (GGTCAAACTCAGC), and a reverse transcriptase (RT) template (TCATCTTTCTCGG). The secondary nicking sgRNA protospacer was GGGTGGGCTCATCTTTCTCG. 200,000 wild-type HEK293 cells were electroporated using the Neon transfection system. After electroporation, cells were clonally isolated by single-cell sorting into 96-well plates using BD FACSAria II. After clonal expansion, genomic DNA was isolated and regions flanking Exon 5 were PCR-amplified using Q5 High-Fidelity DNA Polymerase (NEB) (oligonucleotide primers F: 5’-TTC GTG TCA GAC CCA CTC AGT G-3’ : R: 5’-TTG GCT ATG TTT GGC TGG AAA TC-3’: IDT). Sanger sequencing was employed to detect single-base mutations. Prior to further use, cell lines with the desired mutations were single-cell sorted three times to ensure each cell line represented a homogenous population.

### 4.2 Immunoprecipitations and phosphatase activity assay

Parental, E420K, and E198K variant HEK293 cells were plated and allowed to grow for 48 hours. Cells were lysed by the addition of ice-cold lysis buffer [50 mM Tris-HCl pH 7.4, 150 mM NaCl, 1% Triton X-100, 5 mM EDTA], containing Protease inhibitor Cocktail (ThermoScientific) and passaged several times through a 28G syringe needle. Lysates were clarified by centrifugation at 14,000 x g for 15 minutes. The supernatant was transferred to a new tube, and protein concentrations were determined using RC DC™ Protein Assay (Bio-Rad 5000121). Dynabeads Protein A (100001D) were used to immunoprecipitated (IP) PPP2R5D (ab188323). For the phosphatase assay, beads were incubated with protein lysates for 2 hours at room temperature with rotation, and immunoprecipitated samples were eluted into 50 mM Tris HCl, 150 mM NaCl, pH 7.4. The phosphatase activity assay was performed as described previously (18). Briefly, 20 µL of IPs were dispensed in 1.33x assay buffer [0.15 M NaCl, 30 mM HEPES pH 7.0, 1 mM DTT, 0.1 mg/mL BSA, 1 mM ascorbate, 1 mM MnCl_2_ with a final assay concentration of 75 µM 6,8-difluoro-4-methylumbelliferyl phosphate (DiFMUP) (Invitrogen D6567). Generation of 6,8-difluro-4-methylumbelliferone (DiFMU) was measured with a BioTek Synergy (Ex 360 nm, Em 460 nm). Aliquots of the IPs were eluted with near-boiling 2x SDS-PAGE sample buffer and were used for western analysis to detect total levels of PPP2R5D, which was used to normalize measured activity. To analyze the interaction of the PP2A catalytic subunit, membranes were treated with 0.2 M NaOH for 30 minutes at 30°C before probing for PP2AC subunit (Millipore Sigma 05-421, clone 1D6).

### 4.3 Proliferation Assays

For analysis of growth curves, cells were counted twice before initial plating. HEK293 WT, E198K-het, and E420K-het cells were plated into 60 mm dishes in 3 replicates of 1 x 10^5^ cells. After 24, 48, 72, and 96 hours, detached or dying cells in the media were collected. Cells were then washed once with phosphate buffered saline (PBS), and cells in PBS were added to the tube with cells collected from the media. Adherent cells were detached from the dish with trypsin and combined with the collected cells from the media and PBS. Cells were resuspended and stained with trypan blue for live cell counting. Each plate was counted two times on a TC20 Automated Cell Counter (Bio-Rad 1450102), and the average was recorded for the cell number of that plate.

### 4.4 Western Analysis

Cells were grown to confluent monolayers and lysed by scraping cells in near boiling 2x SDS-PAGE sample buffer (62.5 mM Tris-HCl, pH 6.8, 20% glycerol, 4% SDS, 0.0025% bromophenol blue, and 0.02% ß-mercaptoethanol) followed by mechanical shearing with a 28G syringe needle. Samples were centrifuged at 14,000 x g for 15 minutes, the supernatant was transferred, and protein concentrations were determined using RC DC™ Protein Assay (Bio-Rad 5000121). 25 μg of each protein sample was separated by electrophoresis and proteins were then transferred onto a Trans-Blot® Turbo™ PVDF Membrane (Biorad 10026933) at 2.5 Å for 3 minutes using Trans-Blot® Turbo™ Transfer System (Biorad 1704150). The membranes were blocked at room temperature for 1 hour in Odyssey blocking buffer (LiCor) and then incubated overnight with primary antibodies at 4°C. The next day, membranes were incubated with secondary horseradish peroxidase (HRP) conjugated antibodies for 1 hour at room temperature. Protein bands were visualized with Clarity™ Western ECL Substrate (Bio-Rad 1705060) using Bio-Rad ChemiDoc™ MP Imaging System.

Antibody to PPP2R5D was purchased from Abcam (ab188323). Antibody to PP2AC subunit clone 1D6 (05-421) was purchased from Millipore Sigma. For detection of the PP2A catalytic subunit, membranes were treated with 0.2 M NaOH for 30 minutes at 30 °C before overnight incubation with a primary PP2AC antibody. Primary antibodies to Pan AKT (4691), P-AKT (T308) (13038), P-AKT (S473) (4060), p70 S6 Kinase (34475), P-p70 S6 Kinase (T389) (9205), Ribosomal Protein S6 (2217), P-Ribosomal Protein S6 (S235:236) (2211), and P-Ribosomal Protein S6 (S240:244) (5364) were purchased from Cell Signaling Technology.

Secondary ECL donkey Anti-Rabbit IgG, HRP-linked whole antibodies were purchased from GE Healthcare UK Limited (NA934V), and secondary goat anti-mouse IgG peroxidase-conjugate was purchased from Millipore Sigma (A-5278).

### 4.5 Quantitative proteomic and phosphoproteomic analyses

Cell pellets were resuspended in ice-cold lysis buffer (8 M urea, 25 mM Tris-HCl pH 8.6, 150 mM NaCl, containing phosphatase inhibitors and protease inhibitors (Roche Life Sciences)) and were lysed by sonication. Lysates were subjected to centrifugation (15,000 x g for 30 minutes at 4°C), and supernatants were transferred to a new tube, and the protein concentration was determined using a bicinchoninic acid (BCA) assay (Pierce/ThermoFisher Scientific). DTT and iodoacetamide were added to reduce and alkylate, respectively. Samples were incubated overnight at 37°C with 1:100 (w/w) trypsin. The next day, the trypsin digest was stopped by the addition of 0.25% TFA (final v/v). Precipitated lipids were removed by centrifugation (3500 x g for 15 minutes), and the peptides in the supernatant were desalted over an Oasis HLB 60 mg plate (Waters). An aliquot containing ∼ 100 µg of peptides was removed and labeled with Tandem-Mass-Tag (TMT) reagent (ThermoFisher Scientific). Once labeling efficiency was confirmed to be at least 95%, each reaction was quenched by the addition of hydroxylamine to a final concentration of 0.25% for 10 minutes, mixed, acidified with TFA to a pH of about 2, and desalted over an Oasis HLB 10 mg plate (Waters).

The desalted multiplex was dried by vacuum centrifugation and separated by offline Pentafluorophenyl (PFP)-based reversed-phase HPLC fractionation as previously described (65). TMT-labeled peptides were analyzed on an Orbitrap Lumos mass spectrometer (ThermoScientific) equipped with an Easy-nLC 1200 (ThermoScientific), and raw data was searched and processed as previously described (18). Peptide intensities were adjusted based on total TMT reporter ion intensity in each channel, and log_2_ transformed. P-values were calculated using a two-tailed Student’s t-test, assuming unequal variance.

Phosphopeptide enrichment was achieved using a Fe-NTA phosphopeptide enrichment kit (ThermoFisher) according to instructions provided by the manufacturer and desalted over an Oasis HLB 10 mg plate (Waters). Phosphopeptides were then labeled with TMT reagents, offline separated as described above, and analyzed on the Orbitrap Lumos. The probability of phosphorylation site localization was determined by PhosphoRS (66). Quantification and data analysis were carried out as described above.

### 4.6 Bioinformatics analysis

Motif analysis was performed using singly phosphorylated sites with a phosphorylation localization score ≥ 0.95 that significantly increased or decreased by at least 2-fold after protein correction (67). Upstream-kinase prediction was carried out using KinomeXplorer (26).

### 4.7 Statistical Analysis

GraphPad Prism was used for all statistical analyses and are reported with standard deviation (SD). Unpaired t-tests were performed for analysis between two sample groups or Kruskal-Wallis with Dunn’s multiple comparisons for multiple sample groups and with Dunnett’s for comparison of multiple sample groups compared to WT, as indicated in figure legends. The proliferation assay was analyzed as two-way ANOVA Tukey’s. N is defined as independent biological replicate cell samples, and n is defined as independent technical replicates. Proteomics and phosphoproteomics data were analyzed using Perseus (68, 69).

## ACKNOWLEDGMENTS

pCMV-PE2 was a gift from David Liu (Addgene plasmid # 132775 ; http://n2t.net/addgene:132775; RRID:Addgene_132775). We appreciate obtaining access to the genetic data on SFARI Base.

## DATA AVAILABILITY

Raw MS data for this study are available at MassIVE (MSV000090346) and PRIDE accession (PXD036848). Reviewer password: p1163.

## SUPPORTING INFORMATION

This article contains supporting information.

## Abbreviations

AKT: Protein kinase B
AMPK: AMP-activated protein kinase
ASD: Autism-spectrum disorder
BCA: Bicinchoninic acid
CDK: Cyclin-dependent kinase
CK2: Casein kinase 2
DD: Developmental delay
DiFMU: 6,8-difluro-4-methylumbelliferone
DiFMUP: 6,8-difluoro-4-methylumbelliferyl phosphate
ErbB: EGF receptor family
ERK: Extracellular signal-regulated kinase
GSK3: Glycogen synthase kinase-3
HRP: Horseradish peroxidase
ID: Intellectual disability
IGBP1: Immunoglobulin binding protein 1
IP: Immunoprecipitate
LCMT1: Leucine carboxyl methyltransferase 1
MAPK: Mitogen-activated protein kinase
nCas9: Nicking Cas9
PAK1: Serine/threonine-protein kinase PAK1
PAM: Protospacer adjacent motif
PDPK1: 3-phosphoinositide dependent protein kinase 1
PE3b: Prime Editor 3b
pegRNA: Prime editing guide RNA
PFP: Pentafluorophenyl
PKC: Protein kinase C
PKN: Serine/threonine-protein kinase N
PP2A: Type 2A serine/threonine phosphoprotein phosphatase
PP2AC: Catalytic subunit of PP2A
PPME1: Protein phosphatase methylesterase 1
PPP2R4: Serine/threonine-protein phosphatase 2A regulatory subunit 4
PPP2R5D: Serine/threonine-protein phosphatase 2A 56 kDa regulatory subunit delta isoform
PRKACA: cAMP-dependent protein kinase catalytic subunit alpha
PRKCZ: Protein kinase C zeta type
PTEN: Phosphatidylinositol 3,4,5-triphosphate 3-phosphatase and dual-specificity protein phosphatase PTEN
PTPA: Serine/threonine-protein phosphatase 2A activator (also known as PPP2R4)
RPS6: Ribosomal Protein S6
RSK: Ribosomal S6 kinase (p90S6K)
RT: Reverse transcriptase
S6K: Ribosomal protein S6 kinase
sgRNA: Single guide RNA
TIPRL: TIP41-like protein
TMT: Tandem-Mass-Tag
TSC: Tuberous sclerosis complex

