## Supporting Information for "Quantitative proteomics and phosphoproteomics of PPP2R5D variants reveal deregulation of RPS6 phosphorylation through converging signaling cascades"

Items included:

**Figure S1.** Protein sequence alignment of PPP2R5C (2A5G) and PPP2R5D (2A5D) showing extent of homology.

**Figure S2.** Generation of PPP2R5D E198K-variant cell line.

**Figure S3.** Genomic sequencing of E198K-het variant cell line.

**Figure S4.** Quantitative changes in the proteome and phosphoproteome in PPP2R5D E420K variant cells.

**Figure S5.** Phosphorylation site motif enrichment analysis of sites that were significantly decreased in the E198K and E420K variant cell line.

**Figure S6.** Rapamycin inhibits mTOR signaling in E198K and E420K variant cells.

**Figure S7.** LY2584702 inhibits mTOR signaling in E198K and E420K variant cells.

**Table S1.** Summary of *PPP2R5D* mutations.

Items included as Excel sheets:

**Table S2.** Proteomics and phosphoproteomics data for E198K and E420K PPP2R5D variant cell line analysis.

**Table S3.** PP2A Subunit Abundance as detected by LC-MS/MS.

**Table S4.** Networkin analysis of phosphorylation sites significantly regulated in E198K and E420K variant cell lines.

```
SP|Q13362|2A5G_HUMAN ------------------------------------------------------------
SP|Q14738|2A5D_HUMAN MPYKLKKEKEPPKVAKCTAKPSSSGKDGGGENTEEAQPQPQPQPQPQAQSQPPSSNKRPS 60

SP|Q13362|2A5G_HUMAN --------MLTCNKAGSRMVVDA--------ANSNGPFQPVVLLHIRDVPPADQEKLFIQ 44
SP|Q14738|2A5D_HUMAN NSTPPPTQLSKIKYSGGPQIVKKERRQSSSRFNLSKNRELQKLPALKDSPTQEREELFIQ 120
                             : . : :*.  :*.          * .   :   *  ::* *  ::*:****

SP|Q13362|2A5G_HUMAN KLRQCCVLFDFVSDPLSDLKWKEVKRAALSEMVEYITHNRNVITEPIYPEVVHMFAVNMF 104
SP|Q14738|2A5D_HUMAN KLRQCCVLFDFVSDPLSDLKFKEVKRAGLNEMVEYITHSRDVVTEAIYPEAVTMFSVNLF 180
                     ********************:******.*.********.*:*:** ****.* **:**:*

SP|Q13362|2A5G_HUMAN RTLPPSSNPTGAEFDPEEDEPTLEAAWPHLQLVYEFFLRFLESPDFQPNIAKKYIDQKFV 164
SP|Q14738|2A5D_HUMAN RTLPPSSNPTGAEFDPEEDEPTLEAAWPHLQLVYEFFLRFLESPDFQPNIAKKYIDQKFV 240
                     ************************************************************

SP|Q13362|2A5G_HUMAN LQLLELFDSEDPRERDFLKTTLHRIYGKFLGLRAYIRKQINNIFYRFIYETEHHNGIAEL 224
SP|Q14738|2A5D_HUMAN LALLDLFDSEDPRERDFLKTILHRIYGKFLGLRAYIRRQINHIFYRFIYETEHHNGIAEL 300
                     * **:*************** ****************:***:******************

SP|Q13362|2A5G_HUMAN LEILGSIINGFALPLKEEHKIFLLKVLLPLHKVKSLSVYHPQLAYCVVQFLEKDSTLTEP 284
SP|Q14738|2A5D_HUMAN LEILGSIINGFALPLKEEHKMFLIRVLLPLHKVKSLSVYHPQLAYCVVQFLEKESSLTEP 360
                     ********************:**::****************************:*:****

SP|Q13362|2A5G_HUMAN VVMALLKYWPKTHSPKEVMFLNELEEILDVIEPSEFVKIMEPLFRQLAKCVSSPHFQVAE 344
SP|Q14738|2A5D_HUMAN VIVGLLKFWPKTHSPKEVMFLNELEEILDVIEPSEFSKVMEPLFRQLAKCVSSPHFQVAE 420
                     *::.***:**************************** *:*********************

SP|Q13362|2A5G_HUMAN RALYYWNNEYIMSLISDNAAKILPIMFPSLYRNSKTHWNKTIHGLIYNALKLFMEMNQKL 404
SP|Q14738|2A5D_HUMAN RALYYWNNEYIMSLISDNAARVLPIMFPALYRNSKSHWNKTIHGLIYNALKLFMEMNQKL 480
                     ********************::******:******:************************

SP|Q13362|2A5G_HUMAN FDDCTQQFKAEKLKEKLKMKEREEAWVKIENLAKANPQYTVYSQASTMSIPVAMETDGPL 464
SP|Q14738|2A5D_HUMAN FDDCTQQYKAEKQKGRFRMKEREEMWQKIEELARLNPQYPMFRAPPPLPPVYSMETETPT 540
                     *******:**** * :::****** * ***:**: **** ::     :    :***: *

SP|Q13362|2A5G_HUMAN FEDVQMLRKTVKDEAHQAQKDPKKDRPLARRKSELPQDPHTKKALEAHCRADELASQDGR 524
SP|Q14738|2A5D_HUMAN AEDIQLLKRTVETEAVQMLKDIKKEKVLLRRKSELPQDVYTIKALEAHKRAEEFLTASQE 600
                      **:*:*::**: ** *  ** **:: * ********* :* ****** **:*: : . .

SP|Q13362|2A5G_HUMAN --
SP|Q14738|2A5D_HUMAN AL 602
```

**Figure S1.** Protein sequence alignment of PPP2R5C (2A5G) and PPP2R5D (2A5D) showing extent of homology. Highlighted region (PEEDEPT) corresponds to the acidic loop between alpha-helices 3 and 4 containing E197, E198, and E200.

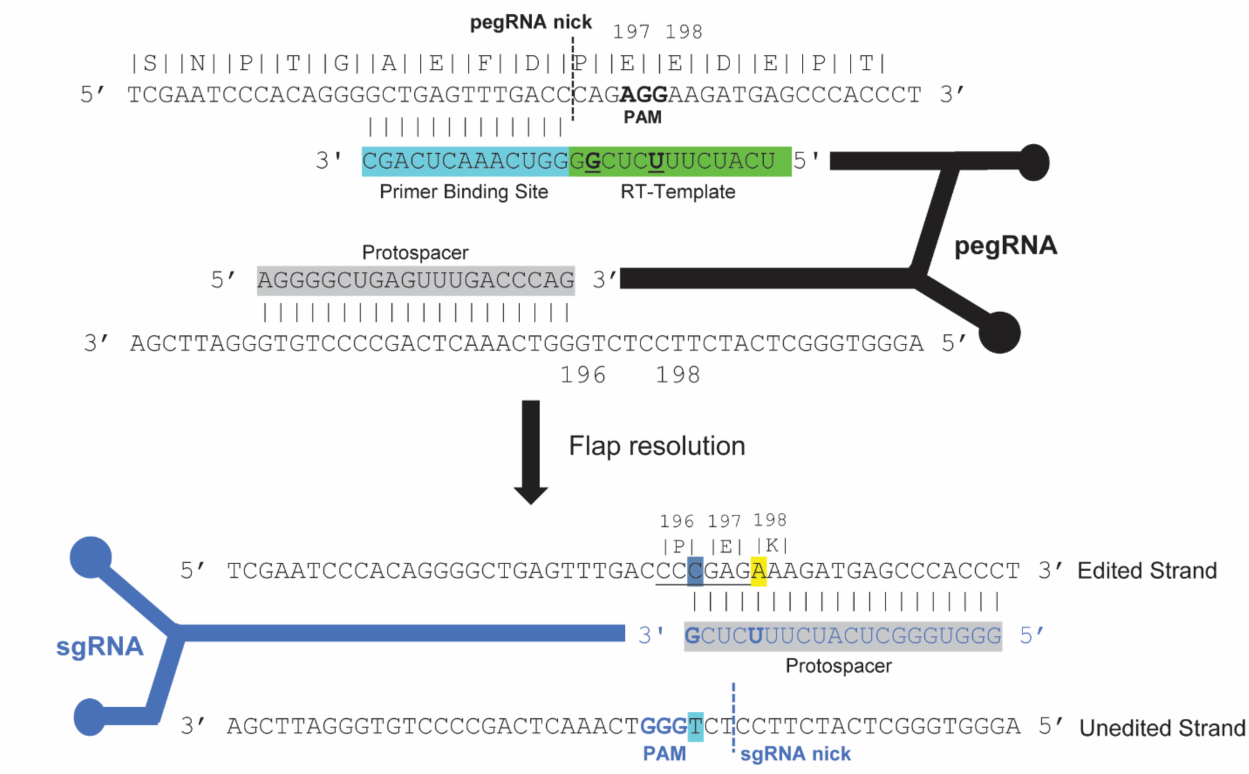


**Figure S2. Generation of PPP2R5D E198K-variant cell line.** Illustration of the prime editing guide RNA (pegRNA) construct used to introduce the nucleotide change c.592G>A into the genome of HEK293 cells via CRISPR-PRIME. The pegRNA was designed to contain the desired edit within the RNA as we as a PAM-silencing mutation to prevent re-occurring nCas9 activity. For targeting c.592G>A, needed to generate the most common variant (E198K) a similar strategy failed, likely due to additional cytosines in close proximity to the desired target. Therefore, to introduce c.592G>A, we used a Prime Editor 3b (PE3b) strategy (Anzalone et al., 2019). The PE3b system is based upon a nicking Cas9 (nCas9) fused to a reverse transcriptase that is programmed by a synthetic prime editing guide RNA (pegRNA), which both targets the specific genomic locus and encodes the desired edit. Using PE3b, the edit occurs without the need for double-strand breaks and a donor template DNA for homology-directed repair. To facilitate prime editing, we designed a pegRNA that positioned the nCas9 to nick the to-be-edited strand upstream of the target site. The PE3b system then allows for the 3'-end of the pegRNA (e.g. the primer binding site; PBS) to hybridize to the 3' end of the nicked strand of genomic DNA. The reverse transcriptase domain then uses the RT template sequence, immediately adjacent to the PBS in the pegRNA, to add complementary deoxyribonucleotides onto the 3' end of the nicked DNA, thus incorporating the desired base change(s) into one DNA strand at the target site. After reverse transcription, a flap resolution nick repair process occurs in which either the newly transcribed 3' DNA sequence is trimmed, and the 5' end of the nicked genomic DNA strand (containing the original WT sequence) is ligated to repair the nick or the WT 5' flap is trimmed and the new 3' sequence, containing the desired edit, is ligated. If the 3' flap resolves, then a base mismatch at the edited site(s) occurs, which could be repaired by the cell using either the WT or edited strand as a template. To bias repair toward the unedited strand (e.g. repair of the unedited strand using the edited strand as a template), a secondary nicking sgRNA is used to target the nCas9-RT prime editor to the DNA strand containing the newly transcribed edit, with subsequent nicking of the unedited strand. In order to reduce the frequency of double-strand breaks and indel formation from potential simultaneous nicking of both DNA strands, the secondary sgRNA is designed to be complementary to the edited sequence and thus contains mismatches to the original WT sequence that should ablate binding to the target site until after the desired edit is resolved. To enhance this effect, a second synonymous mutation of proline 196 (CCA= Pro to CCC= Pro) was included in the RT template. This produces two PAM-proximal seed region base mismatches for sgRNA binding to the WT sequence, which can dramatically reduce secondary nicking of unedited DNA. Additionally, the introduction of the E198K mutation destroys the AGG PAM utilized by the pegRNA, thereby reducing or preventing Prime Editor activity at the site once the desired edit has been fully resolved. When the strategy of introducing a secondary synonymous mutation to suppress DSBs is adopted, it is important to make sure that the codon usage for the synonymous change (e.g. CCA= Pro; CCC= Pro) is similar to the WT codon, in order to guard against introducing an extraneous genomic change that could alter the efficiency of translation. After electroporation with the PE3b system components, the cells were grown for 48 hours to ensure cell replication occurred, which makes the mutation permanent in the genome.

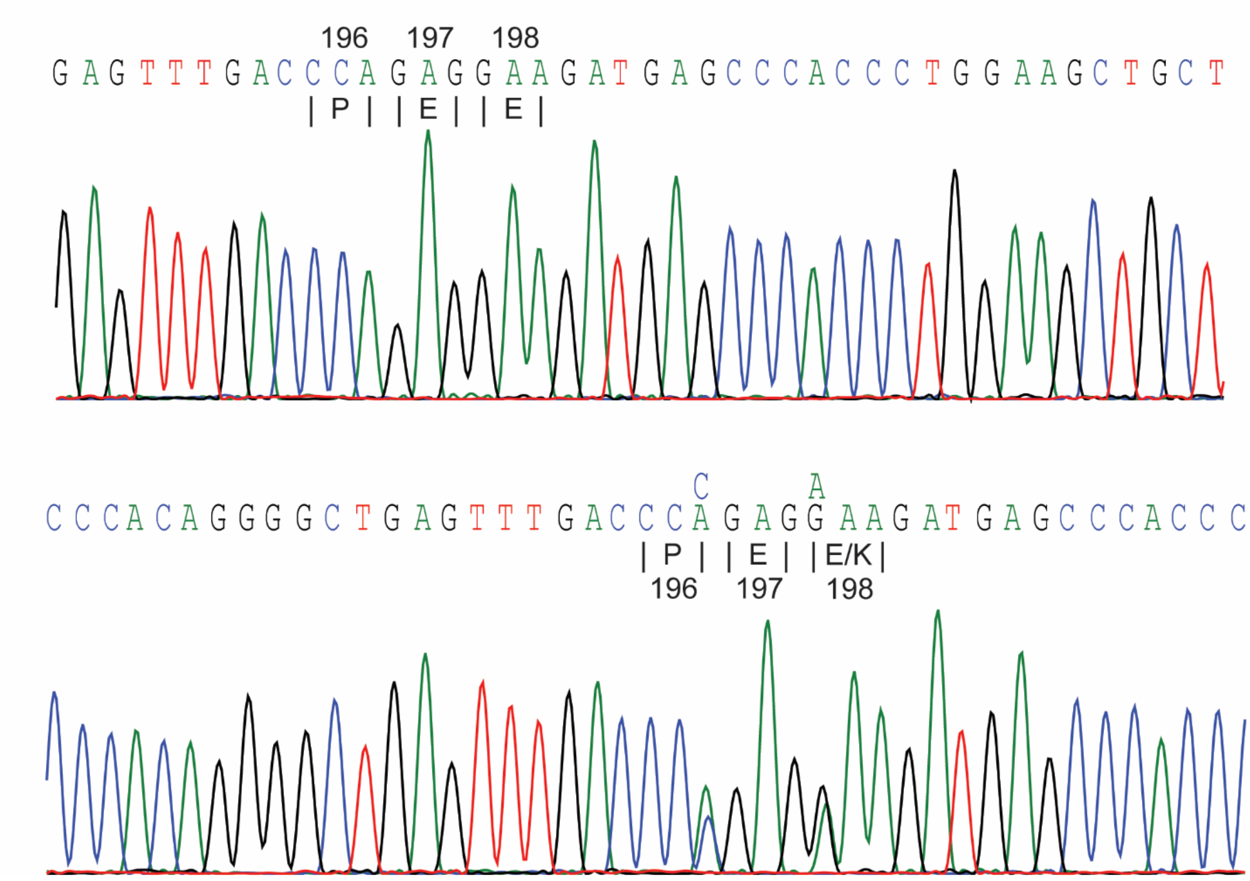


**Figure S3. Genomic sequencing of E198K-het variant cell line.** Sanger sequencing data from WT encoding glutamic acid (GAA) in both alleles and E198K heterozygous (E198K-het) encoding glutamic acid (GAA) in one allele and lysine (AAA) in the other.

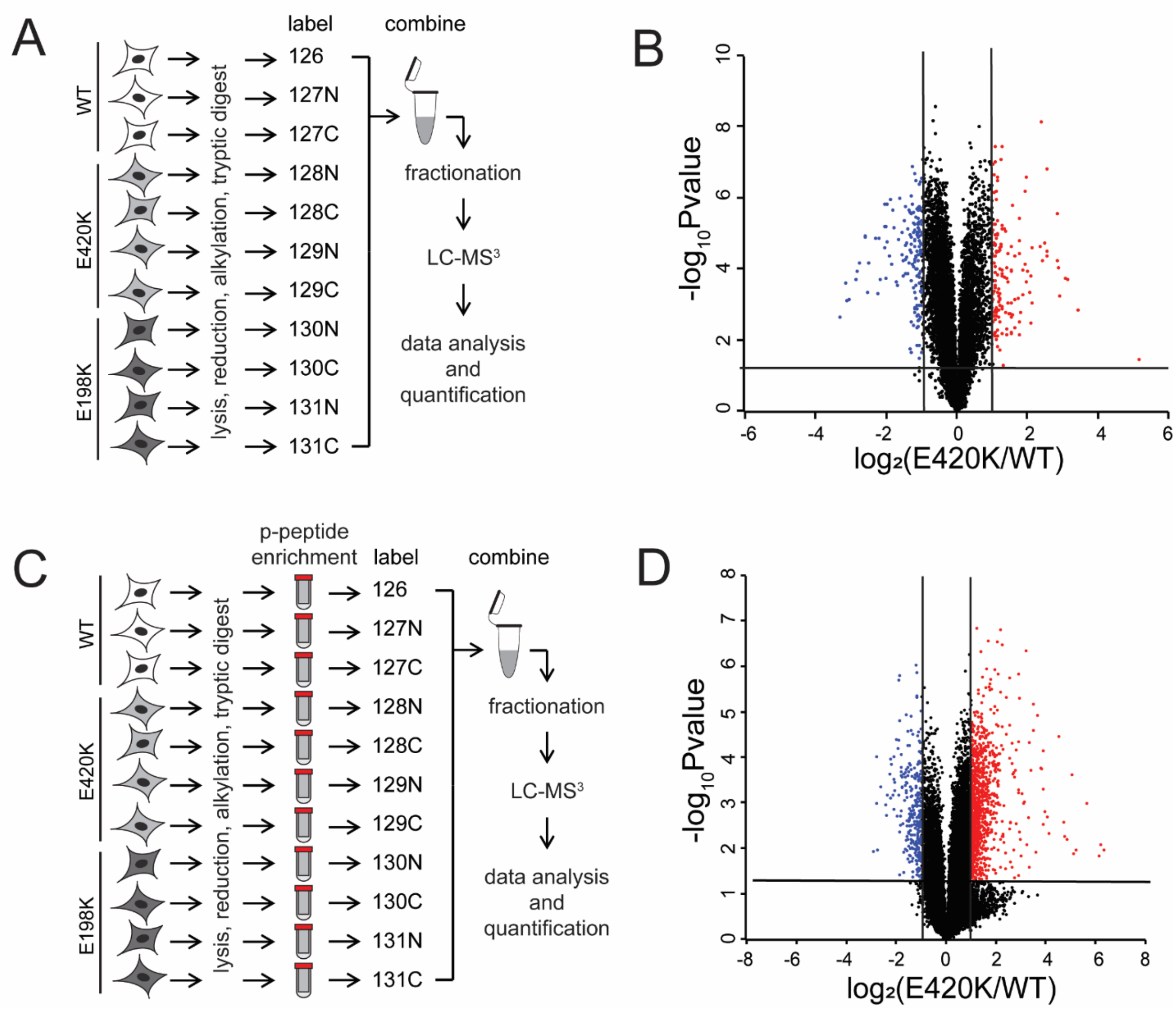


**Figure S4. Quantitative changes in the proteome and phosphoproteome in PPP2R5D E420K variant cells.** Workflow for the quantitative proteomics (A) and phosphoproteomics (C) protocol used. Volcano plots for proteomic (B) and protein-corrected phosphoproteomic changes (D) in E420K variant cells (N=3 and N=4 independent biological replicates for WT and E420K-het, respectively.) Volcano plots show $\log_2$ fold change versus the negative $\log_{10}$ of the *p*-value of the fold change. Statistical significance corresponding to a *p*-value of <0.05 is shown by peptides with a $-\log_{10}$ value of 1.3 or greater. Peptides shown in blue or red are two-fold or more decreased or increased in abundance, respectively.

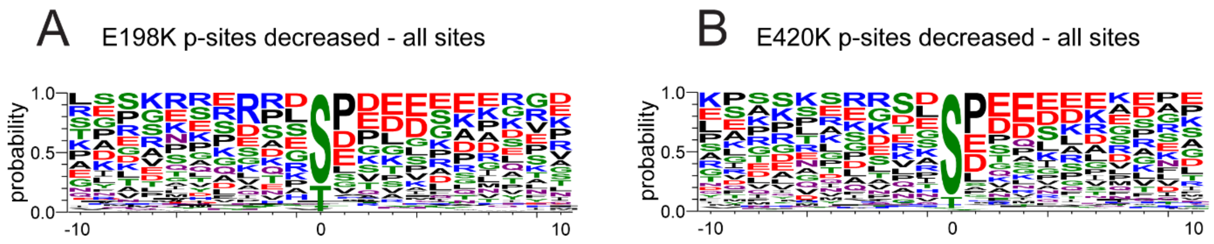


**Figure S5. Phosphorylation site motif enrichment analysis of sites that were significantly decreased in the E198K and E420K variant cell line.** (A) Weblogos showing enrichment of amino acids surrounding the phosphorylated residue for sites significantly decreased by at least two-fold in PPP2R5D E198K (A) and E420K (B) variant cells.

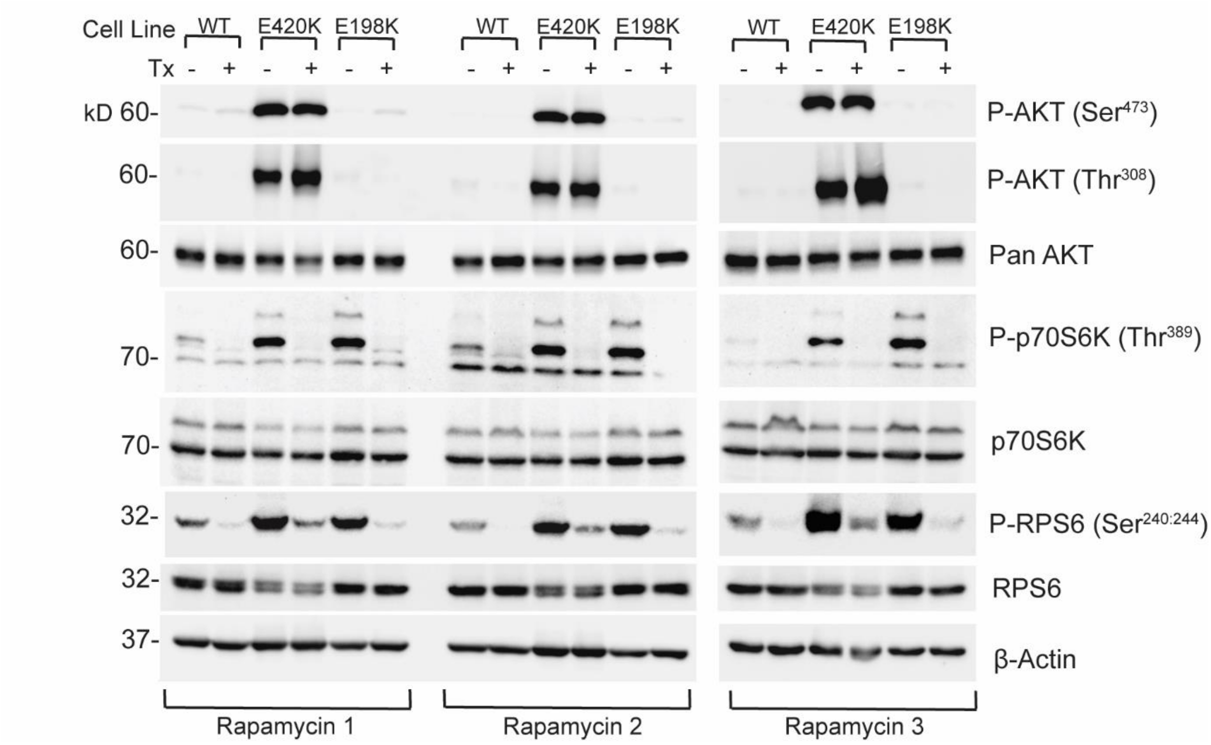


**Figure S6. Rapamycin inhibits mTOR signaling in E198K and E420K variant cells.** Immunoblots of WT, PPP2R5D E420K-het, and PPP2R5D E198K-het HEK293 cell lines treated with rapamycin (10nm for 1 hour). Treatment was performed in three independent experiments.

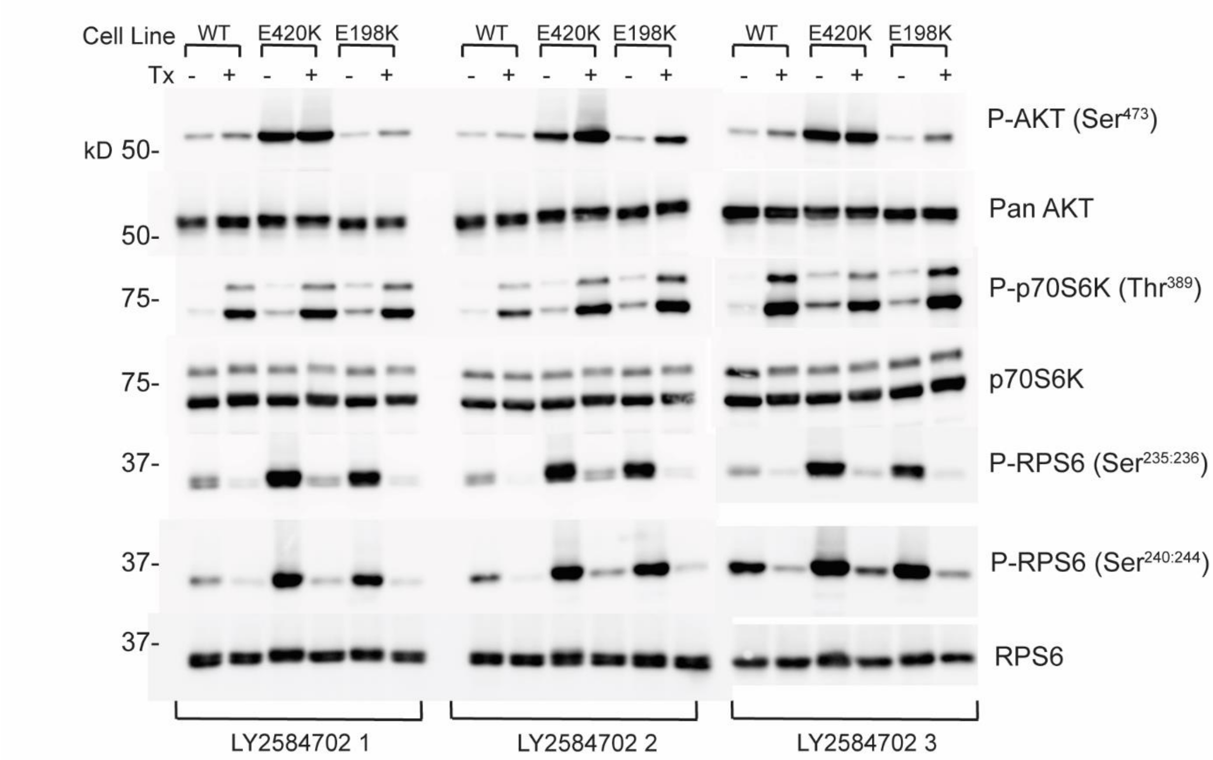


**Figure S7. LY2584702 inhibits mTOR signaling in E198K and E420K variant cells.** Immunoblots of WT, PPP2R5D E420K-het, and PPP2R5D E198K-het HEK293 cell lines treated with LY2584702 (20 nM for 3 hours). Treatment was performed in three independent experiments.

| *PPP2R5D* Pathogenic Variant | | # cases in ClinVar | # cases in SFARI |
|---|---|---|---|
| E34A | | 1 | - |
| P53S | | 2 | - |
| S54P | | 1 | - |
| R85* | | 1 | 1 |
| E197K | | 3 | 2 |
| E197G | | 1 | 6 |
| E198K | | 22 | 34 |
| E200K | | 15 | 11 |
| P201R | | 3 | - |
| Glu200_Pro201delinsGlyHis | | 1 | 1 |
| W207R | | 4 | 5 |
| Q211P | | 4 | 2 |
| R219C | | 1 | - |
| I230T | | 1 | 1 |
| E250K | | 3 | |
| D251A | | 3 | 3 |
| D251H | | 1 | 1 |
| D251V | | 3 | 4 |
| D251Y | | 3 | 2 |
| R253G | | 1 | - |
| R253P | | 2 | 2 |
| R253Q | | 3 | - |
| H263R | | 1 | - |
| L313V | | 2 | 1 |
| E420K | | 5 | 4 |
| F473C | | 1 | - |
| F473L | | 2 | 1 |
| P525L | | 1 | - |
| E537K | | 1 | - |
| H588Y | | 1 | - |

**Table S1. Summary of *PPP2R5D* mutations.** Number of cases of pathogenic PPP2R5D variants reported in NCBI’s ClinVar and Simons Foundation Autism Research Initiative (SFARI).
